# Targeting eIF2α in TBI-induced traumatic optic neuropathy: Effects of Salubrinal and the Integrated Stress Response Inhibitor

**DOI:** 10.1101/2024.02.15.580435

**Authors:** Shelby M. Hetzer, Rohan Bellary, Jordyn N. Torrens, Robert F. Grimaldi, Nathan K. Evanson

## Abstract

Traumatic brain injury (TBI) can induce traumatic axonal injury in the optic nerve, which is referred to as traumatic optic neuropathy (TON). TON occurs in up to 5% of TBI cases and leads to irreversible visual deficits. TON-induced phosphorylation of eIF2α, a downstream ER stress activator in the PERK pathway presents a potential point for therapeutic intervention. For eIF2α phosphorylation can lead to apoptosis or adaptation to stress. We hypothesized that dephosphorylation, rather than phosphorylation, of eIF2α would lead to reduced apoptosis and improved visual performance and retinal cell survival. Adult male mice were injected with Salubrinal (increases p-eIF2α) or ISRIB (decreases p-eIF2α) 60 minutes post-injury. Contrary to literature, both drugs hindered control animal visual function with minimal improvements in injured mice. Additionally, differences in eIF2α phosphorylation, antioxidant responses, and protein folding chaperones were different when examining protein expression between the retina and its axons in the optic nerve. These results reveal important compartmentalized ER stress responses to axon injury and suggest that interventions in the PERK pathway may alter necessary homeostatic regulation of the UPR in the retina.

## Introduction

Traumatic brain injury (TBI) affects approximately 2.9 million people yearly (Taylor CA, 2017). TBI first induces a primary injury that manifests directly then secondary effects arise that tend to cause long-term complications due to molecular signaling cascades, leading to chronic neurological complications (Ladak et al., 2019). Moreover, the results of TBI are widespread and not limited to any one brain-controlled system. This variability or lack of regional specificity results from the inherent diversity of TBIs where no one is identical to another. Importantly, many of the widespread, divergent symptomologies are likely driven by diffuse axonal injury which has an estimated incidence of 73% of all TBI cases.

One way that axonal injury impacts TBI patients can be seen through the array of visual defects that arise in 50-60% of patients (Ventura et al., 2014), with 2-3% attributed to injury of the optic nerve (Chen et al., 2022). This optic nerve axonal trauma, or traumatic optic neuropathy (TON), is one pathophysiological outcome associated with TBI (Steinsapir and Goldberg, 2011). As a result of injury to the optic nerve, patients experience a range of symptoms that include progressive retinal thinning, visual acuity decreases, or outright vision loss. Some longitudinal case studies report progressive visual changes as far after injury as 30 years (Chen et al., 2017; Chan et al., 2019). Due to the concern surrounding this condition and the lack of knowledge regarding the mechanisms, treatment, or chronic consequences, there is a clear need to discover potential interventions (Bastakis et al., 2019; Wladis et al., 2021; Karimi et al., 2021).

Nevertheless, TON remains understudied in this context despite emerging evidence that it is likely a more prevalent co-morbidity of TBI than previously assumed (Evans et al., 2021; Hetzer et al., 2023). Our weight-drop TBI model provides an opportune system for exploring the effects of traumatic axonal injury to the optic nerve as we have shown that our closed-head injury produces replicable damage to the optic nerve with subsequent death of retinal ganglion cells (RGCs). We also previously found that TBI-induced TON leads to elevation of endoplasmic reticulum (ER) stress markers in the retina post-injury (Hetzer et al., 2021; Torrens et al., 2023). Of the three ER stress receptors activated under duress (IRE1α, PERK, and/or ATF6), both the inositol-requiring enzyme 1 alpha (IRE1α) and PERK pathways were elevated up to 30 days post-injury. However, the PERK pathway was more sensitive to chronic upregulation and a brief-oxygen intervention in our model (Torrens et al., 2023), so we decided to intervene in this pathway to assess effects on visual outcomes. There are two readily available pharmaceuticals previously tested in the context of TBI – Salubrinal (Rubovitch et al., 2015; Tan et al., 2018; Wang et al., 2019; Logsdon et al., 2016) and the Integrated Stress Response Inhibitor (ISRIB) (Chou et al., 2017; Wenzhu Zhou et al., 2023). Both drugs target activity of PERK’s downstream eukaryotic translation initiation factor, eIF2α.

Once phosphorylated, following the heterodimerization of PERK, the alpha subunit of the tertiary eIF2 complex can act to improve ER stress by halting protein translation and/or promoting transcription of downstream effector mRNAs like antioxidants and apoptotic factors like CHOP (C/EBP Homologous Protein). Phosphorylated eIF2α (p-eIF2α) inhibits binding of its guanine nucleotide exchange factor, eIF2B (Konieczny and Safer, 1983). Without eIF2B, eIF2α cannot be returned to its active GTP-bound state, thus halting transcription due to a lack of eIF2’s complete ternary complex (Bogorad et al., 2018). Interestingly, depending on the severity of the stress in the ER, eIF2α can lead to translation of open reading frames for proteins that can assist with both cell survival (e.g., antioxidants, metabolic support, nutrient uptake, mitochondrial function) and cell death (e.g., CHOP, Caspase 12, autophagy genes (Bond et al., 2020)). eIF2α, however, is not solely associated with ER stress, but is also the crux of the Integrated Stress Response, which can respond to oxidative stress, heat shock, DNA damage, hypoxia, amino acid deprivation, viral infection, and ultraviolet light (Wek, 2018; Bond et al., 2020). Upregulation of PERK-phosphorylation with eIF2α, along with IRE1α pathway activation, indicates ER stress, which is what we have shown in our model.

Given that the PERK pathway is upregulated in TON-induced visual deficits and sustained retinal cellular loss post-TBI (Torrens et al., 2023; Hetzer et al., 2021), the PERK-eIF2α pathway poses a potential control point in preventing the secondary cellular outcomes of TON in retinal cells. From this duality of eIF2α-based stress control, we chose to compare effects of contrasting drugs, Salubrinal and ISRIB. Salubrinal promotes the continued phosphorylation of eIF2α to maintain decreased protein translation and reduce the burden on the ER during injured or diseased states. In contrast, ISRIB is a small-molecule inhibitor that prevents the formation of stress granules impairing eIF2α’s ability to translate downstream factors like the pro-apoptotic Activating transcription factor 4 (ATF4)-CHOP cascade (Sidrauski et al., 2015). ISRIB appears to only induce this downregulation of mRNA translation in stressed cells, leaving unstressed cells unaltered (Sidrauski et al., 2015). As noted above, both of these drugs show promise in treating experimental TBI but neither has been tested in the context of TON or other models of optic nerve injury.

We hypothesized that manipulating the PERK pathway via ISRIB-mediated dephosphorylation of eIF2α would lead to increased retinal ganglion cell survival and improved visual function in a TBI-induced TON mouse model. Conversely, we predict opposite effects with Salubrinal, which we hypothesize will lead to increased apoptotic factors to induce ER-mediated cell death of RGCs (Figure 1).

**Figure 1.**
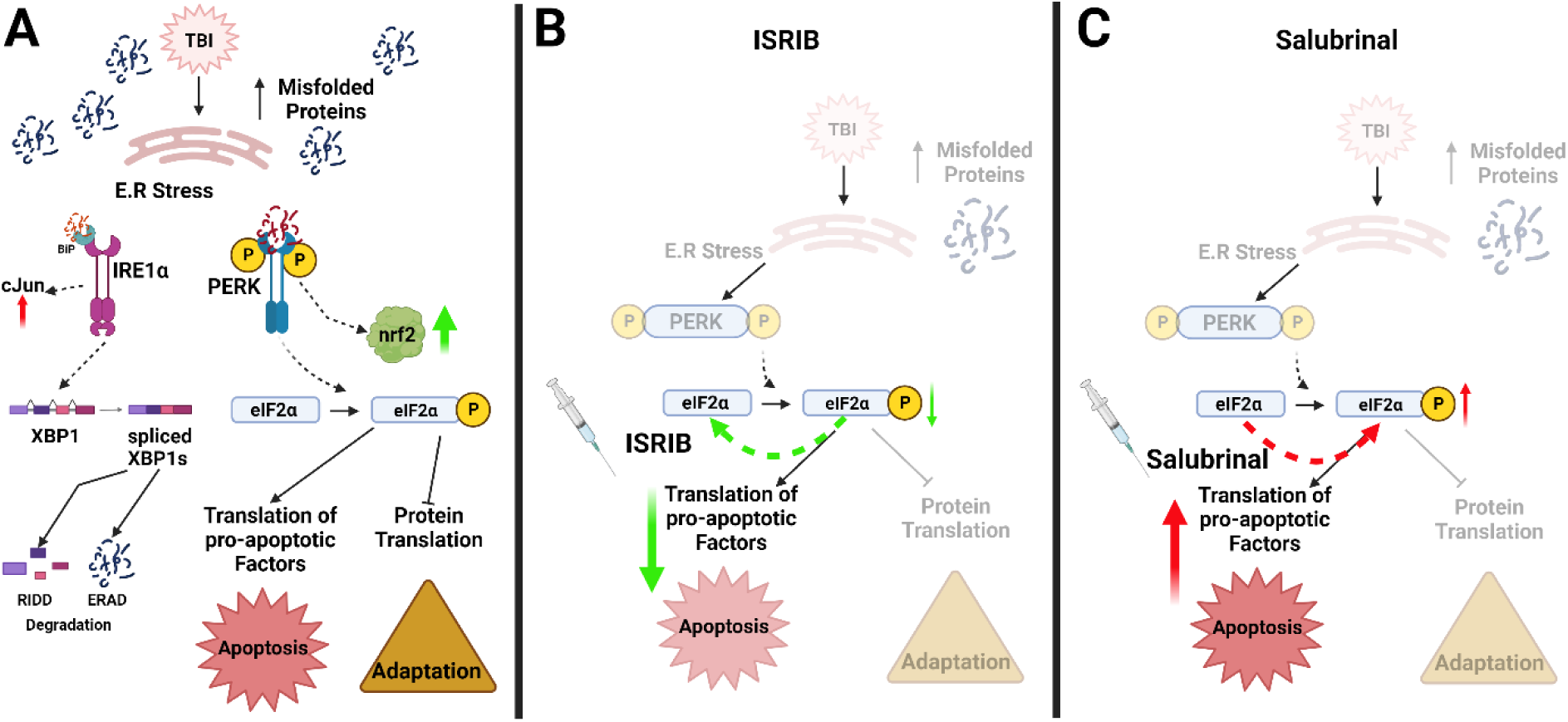
Predicted effects of Salubrinal and ISRIB. **(A)** Depicts the basic pathways for ER stress-mediated activation of IRE1α and PERK pathways. Upon binding of misfolded proteins to IRE1α or PERK (or their pre-bound Binding Immunoglobulin Protein, BiP), these receptors can initiate cascades for both adaptation to injury by reduction of ER burden via mRNA degradation and protein translation inhibition or apoptosis. IRE1α will splice XBP1 allowing it to initiate mRNA degradation (regulated IRE1α-dependent mRNA decay [RIDD]) or tag proteins for exportation and degradation by the proteosome (ER associated degradation [ERAD]). Upon too great or prolonged ER stress, IRE1α can also activate the pro-apoptotic cJun pathway. PERK phosphorylates eIF2α, which is able to halt protein translation (adaptive to an extent) while also facilitating translation of a select few factors like ATF4, which can then go on to the nucleus to initiate translation of apoptotic genes (e.g., CHOP) autophagy genes, biosynthetic genes, and antioxidant genes (e.g., nrf2). **(B)** We hypothesize that ISRIB-mediated dephosphorylation of eIF2α will be most beneficial by preventing the upregulation of proteins like ATF4 and CHOP as well as potential feedback for inactivation of the IRE1α pathway. **(C)** In contrast, we hypothesize that increasing eIF2α phosphorylation with Salubrinal will lead to worse outcomes through increased apoptotic protein translation.

## Methods

### Animals and Interventions

Eight-week-old, male mice (C57BI/6J) underwent a closed-head weight drop TBI as previously described (Evanson et al., 2018). TBI occurred on day 0 of experimental procedures (Figure 2c). SHAM mice were anesthetized but did not undergo TBI. Mice were then injected intraperitoneally one-hour post-injury with a single dose of Salubrinal (1.5mg/kg), ISRIB (2.5 mg/kg), or Vehicle (6.25% DMSO in sterile saline). Doses were chosen based on previously published research to be sufficient to produce beneficial effects (Chou et al., 2017; Wang et al., 2019) and one-hour post injury dosing was chosen because preliminary data suggested that ER stress begins to elevate one hour post injury and remains elevated until at least 3 hours post injury. Mice were thus split into six cohorts – Salubrinal (n=24), ISRIB (n= 24), or Vehicle (n= 24) intervention plus or minus TBI (TBI n = 36, SHAM n=36). Supplementary Figure 1 shows a detailed timeline of experimental procedures.

**Figure 2.**
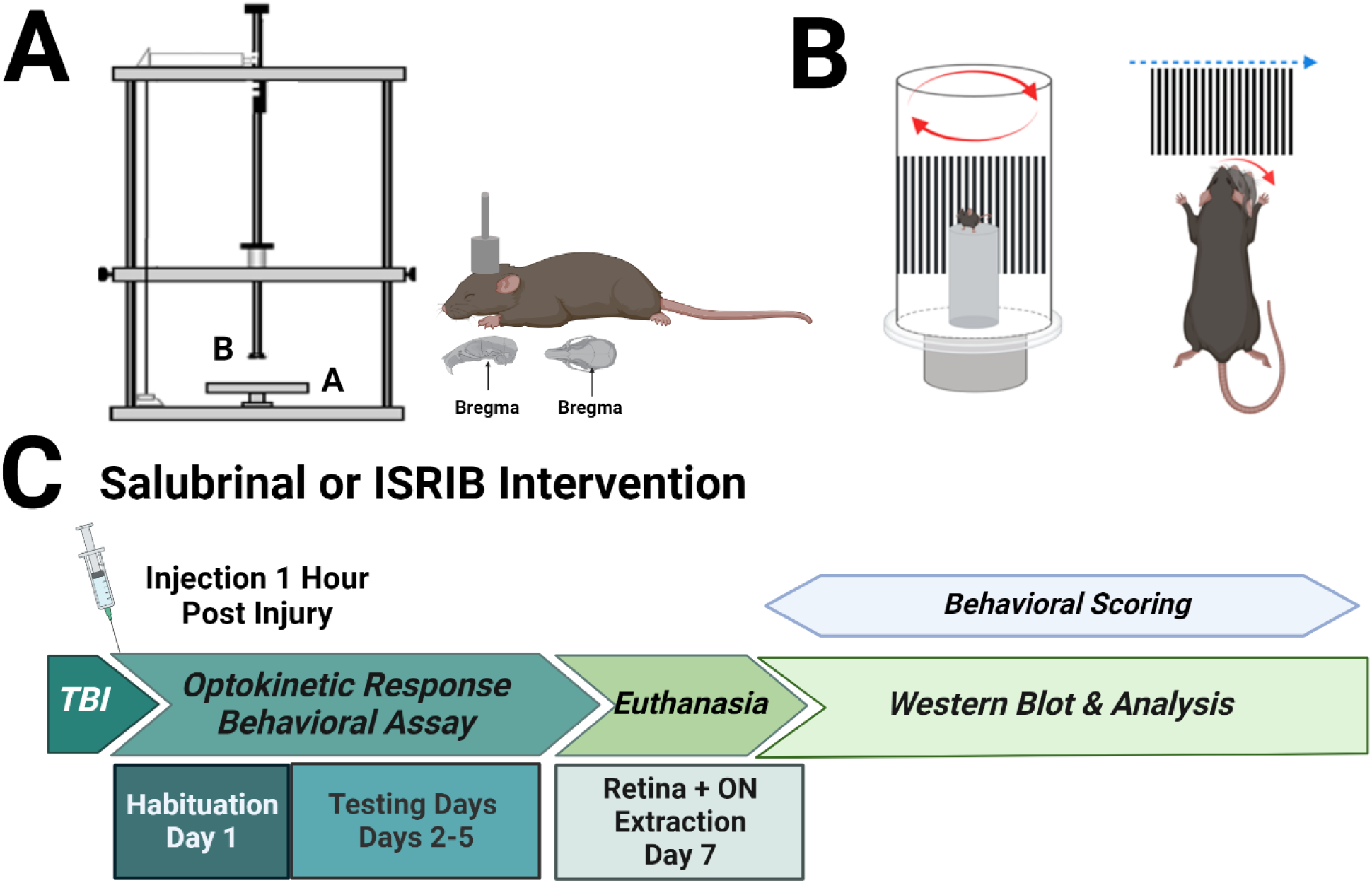
Experimental Timeline and Procedures. In **(A)** we show a depictions of our weight drop device along with the positioning of the weight atop the head. Within **A** the smaller “A” represents the platform and “B” the weight. We also show a depiction of our optomotor device **(B)** and our timeline of procedures **(C)**.

### Optomotor Response (OMR) Testing

As previously described (Hetzer et al., 2021), mice underwent four days of behavioral testing for optomotor nystagmus function using a custom built optomotor behavioral assay (Figure 2b). Briefly, optomotor testing utilizes the manipulation of sine-wave gratings ranging from wider to thinner alternating black and white bars to assess involuntary visual responses in mice (0.12, 0.26, 0.32, and 0.39 cycles per degree [cpd]). Mice are placed on an immobile platform as the gratings rotate around them in both clockwise and counterclockwise direction at two rpm for two minutes in each direction. A visual response in this task is the involuntary visual tracking of the moving stimulus, termed an optomotor response (OMR). Thus, this device can be used to assess the basic functionality of the optomotor response (i.e., is there a response or is it blunted) in addition to visual acuity. Visual acuity is the ability of a subject to detect distinct visual stimuli (i.e., distinguishing the white and black bars as separate). This was assessed by the varying the cycles/degree to determine thresholds of the OMR. These gratings are shown on the x-axis of Figure 3. Higher cpds were indicative of thinner bars that are more “difficult” for the subject to discern, thereby providing us with information on the threshold of visual acuity, the frequency at which OMRs stop (Thaung et al., 2002). Statistics are presented in Table 1.

**Figure 3.**
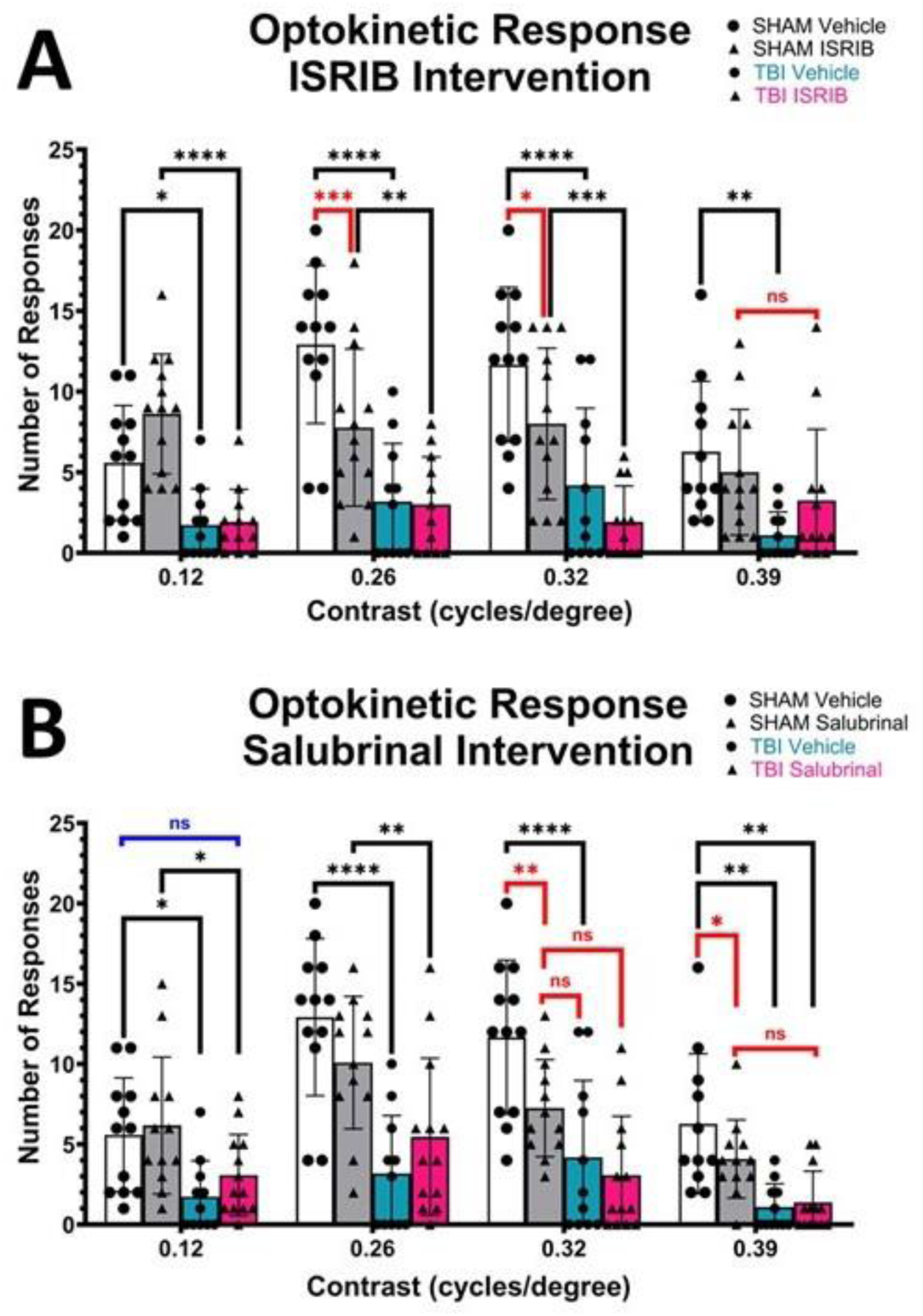
Optomotor Function. **(A)** ISRIB did not improve optomotor performance within the first 7 days post injury. Instead, we show that ISRIB may have been more detrimental to retinal cell function as SHAM ISRIB mice performed worse than vehicle controls. Still, ISRIB may better preserve visual acuity as TBI ISRIB mice performed similarly to controls at 0.39cpd. **(B)** Salubrinal results are largely the same, suggesting that there are inherent/homeostatic needs in the retina for PERK pathway function. Blue text highlights comparisons showing either benefits of interventions between sham and injured mice while red text highlights potential negative effects in sham mice due to intervention. Data was analyzed by 2-way ANOVA (individual analysis for each contrast; injury x drug) and is presented as mean ± SEM with each symbol representing an individual animal. n=12/group; * p<0.05, ** p<0.01, *** p<0.001, **** P<0.0001

**Table 1.** OMR Statistics.

| SALUBRINAL |  |  |  |  |  |  |  |  |
| --- | --- | --- | --- | --- | --- | --- | --- | --- |
| Grating 0.12 |  |  |  |  |  |  |  |  |
|  | Test Statistic | p value | Post Hoc Comparrison's p values |  |  |  |  |  |
| Injury | F (1,44) = 13.63 | 0.006 | SHAM Veh v TBI Veh | SHAM Veh vs SHAM Sal | SHAM Veh vs TBI Sal | TBI Veh vs SHAM Sal | TBI Veh vs TBI Sal | SHAM Salvs TBI Sal |
| Drug | F(1,44)=1.05 | 0.31 | 0.007 | 0.66 | 0.06 | 0.002 | 0.32 | 0.02 |
| Interaction | F(1,44)=0.16 | 0.69 |  |  |  |  |  |  |
| Grating 0.26 |  |  |  |  |  |  |  |  |
|  | Test Statistic | p value | Post Hoc Comparrison's p values |  |  |  |  |  |
| Injury | F (1,44) = 3.96 | <0.0001 | SHAM Veh v TBI Veh | SHAM Veh vs SHAM Sal | SHAM Veh vs TBI Sal | TBI Veh vs SHAM Sal | TBI Veh vs TBI Sal | SHAM Salvs TBI Sal |
| Drug | F(1,44)=0.04 | 0.83 | <0.0001 | 0.12 | 0.0001 | 0.0006 | 0.21 | 0.01 |
| Interaction | F(1,44)=0.16 | 0.05 |  |  |  |  |  |  |
| Grating 0.32 |  |  |  |  |  |  |  |  |
|  | Test Statistic | p value | Post Hoc Comparrison's p values |  |  |  |  |  |
| Injury | F (1,44) = 24.05 | <0.0001 | SHAM Veh v TBI Veh | SHAM Veh vs SHAM Sal | SHAM Veh vs TBI Sal | TBI Veh vs SHAM Sal | TBI Veh vs TBI Sal | SHAM Salvs TBI Sal |
| Drug | F(1,44)=5.4 | 0.02 | 0.0004 | 0.06 | <0.0001 | 0.29 | 0.91 | 0.07 |
| Interaction | F(1,44)=0.19 | 0.17 |  |  |  |  |  |  |
| Grating 0.39 |  |  |  |  |  |  |  |  |
|  | Test Statistic | p value | Post Hoc Comparrison's p values |  |  |  |  |  |
| Injury | F (1,44) = 6.81 | <0.0001 | SHAM Veh v TBI Veh | SHAM Veh vs SHAM Sal | SHAM Veh vs TBI Sal | TBI Veh vs SHAM Sal | TBI Veh vs TBI Sal | SHAM Salvs TBI Sal |
| Drug | F(1,44)=0.04 | 0.04 | <0.0001 | 0.008 | <0.0001 | 0.02 | 0.99 | 0.03 |
| **Interaction** | F(1,44)=6.81 | **0.01** |  |  |  |  |  |  |
| **ISRIB** |  |  |  |  |  |  |  |  |
| **Grating 0.12** |  |  |  |  |  |  |  |  |
|  | **Test Statistic** | **p value** | **Post Hoc Comparrisonns p values** |  |  |  |  |  |
| **Injury** | F (1,44) = 36.72 | **<0.0001** | **SHAM Veh v TBI Veh** | **SHAM Veh vs SHAM ISRIB** | **SHAM Veh vs TBI ISRIB** | **TBI Veh vs SHAM ISRIB** | **TBI Veh vs TBI ISRIB** | **SHAM ISRIBvs TBI ISRIB** |
| **Drug** | F(1,44)=3.42 | 0.07 | **0.02** | 0.07 | **0.02** | **<0.0001** | 0.99 | **<0.0001** |
| **Interaction** | F(1,44)=2.66 | 0.11 |  |  |  |  |  |  |
| **Grating 0.26** |  |  |  |  |  |  |  |  |
|  | **Test Statistic** | **p value** | **Post Hoc Comparrisonns p values** |  |  |  |  |  |
| **Injury** | F (1,44) = 35.80 | **<0.0001** | **SHAM Veh v TBI Veh** | **SHAM Veh vs SHAM ISRIB** | **SHAM Veh vs TBI ISRIB** | **TBI Veh vs SHAM ISRIB** | **TBI Veh vs TBI ISRIB** | **SHAM ISRIBvs TBI ISRIB** |
| **Drug** | F(1,44)=4.83 | **0.03** | **<0.0001** | **0.02** | **<0.0001** | **0.04** | 0.99 | **0.03** |
| **Interaction** | F(1,44)=4.2 | **0.04** |  |  |  |  |  |  |
| **Grating 0.32** |  |  |  |  |  |  |  |  |
|  | **Test Statistic** | **p value** | **Post Hoc Comparrisonns p values** |  |  |  |  |  |
| **Injury** | F (1,44) = 30.22 | **<0.0001** | **SHAM Veh v TBI Veh** | **SHAM Veh vs SHAM ISRIB** | **SHAM Veh vs TBI ISRIB** | **TBI Veh vs SHAM ISRIB** | **TBI Veh vs TBI ISRIB** | **SHAM ISRIBvs TBI ISRIB** |
| **Drug** | F(1,44)=5.77 | **0.02** | **0.0007** | 0.15 | **<0.0001** | 0.14 | 0.58 | **0.005** |
| **Interaction** | F(1,44)=0.32 | 0.57 |  |  |  |  |  |  |
| **Grating 0.39** |  |  |  |  |  |  |  |  |
|  | **Test Statistic** | **p value** | **Post Hoc Comparrisonns p values** |  |  |  |  |  |
| **Injury** | F (1,44) = 15.9 | **0.0002** | **SHAM Veh v TBI Veh** | **SHAM Veh vs SHAM ISRIB** | **SHAM Veh vs TBI ISRIB** | **TBI Veh vs SHAM ISRIB** | **TBI Veh vs TBI ISRIB** | **SHAM ISRIBvs TBI ISRIB** |
| **Drug** | F(1,44)=0.002 | 0.96 | **0.0005** | 0.37 | **0.03** | **0.03** | 0.44 | 0.58 |
| **Interaction** | F(1,44)=0.03 | **0.03** |  |  |  |  |  |  |

### Western Blotting

Mice were euthanized by rapid decapitation seven days post injury, and retinas and optic nerves were extracted into lysis buffer (RIPA Buffer, Sigma Aldrich, CAT: R0278) plus Halt protease and phosphatase inhibitor (Thermo Fisher, CAT: 78440) and frozen on dry ice. Retinas and optic nerves were then lysed, and a BCA assay was used to determine protein concentrations and subsequent loading amounts (20-30µg). Controls of a sham brain (intermembrane) and water (negative) were used on each blot. Twenty µg retinal samples and 15µg nerve samples were loaded onto SDS-PAGE gels and transferred onto Amersham Hybond-P 0.45 µm PVDF membranes (GE Life Sciences, Pittsburgh, PA; CAT: GE 10600029). Membranes were incubated in Fisher No-Stain Total Protein Stain (Thermo Fisher, CAT: A44449) as per manufacturer instructions, blocked in either 5% non-fat milk or 5% BSA (manufacturer dependent) for 1 hour at room temperature, followed by overnight incubation in primary antibodies at 4°C (see antibody table 2). Membranes were rinsed in TBST then incubated in anti-rabbit HRP for two hours at room temperature. Blots were imaged using an iBright™ Imaging System (Thermo Fisher) and analyzed using Image J. Some blots were stripped following imaging in stripping buffer (99% β-mercaptoethanol, 20% Sodium Dodecyl Sulfate, and 1M Tris-HCl pH 6.8) for 30 minutes at 37°C, washed, re-blocked, and exposed to the same immunoblotting steps as above. Statistics are presented in Table 3.

**Table 2.**
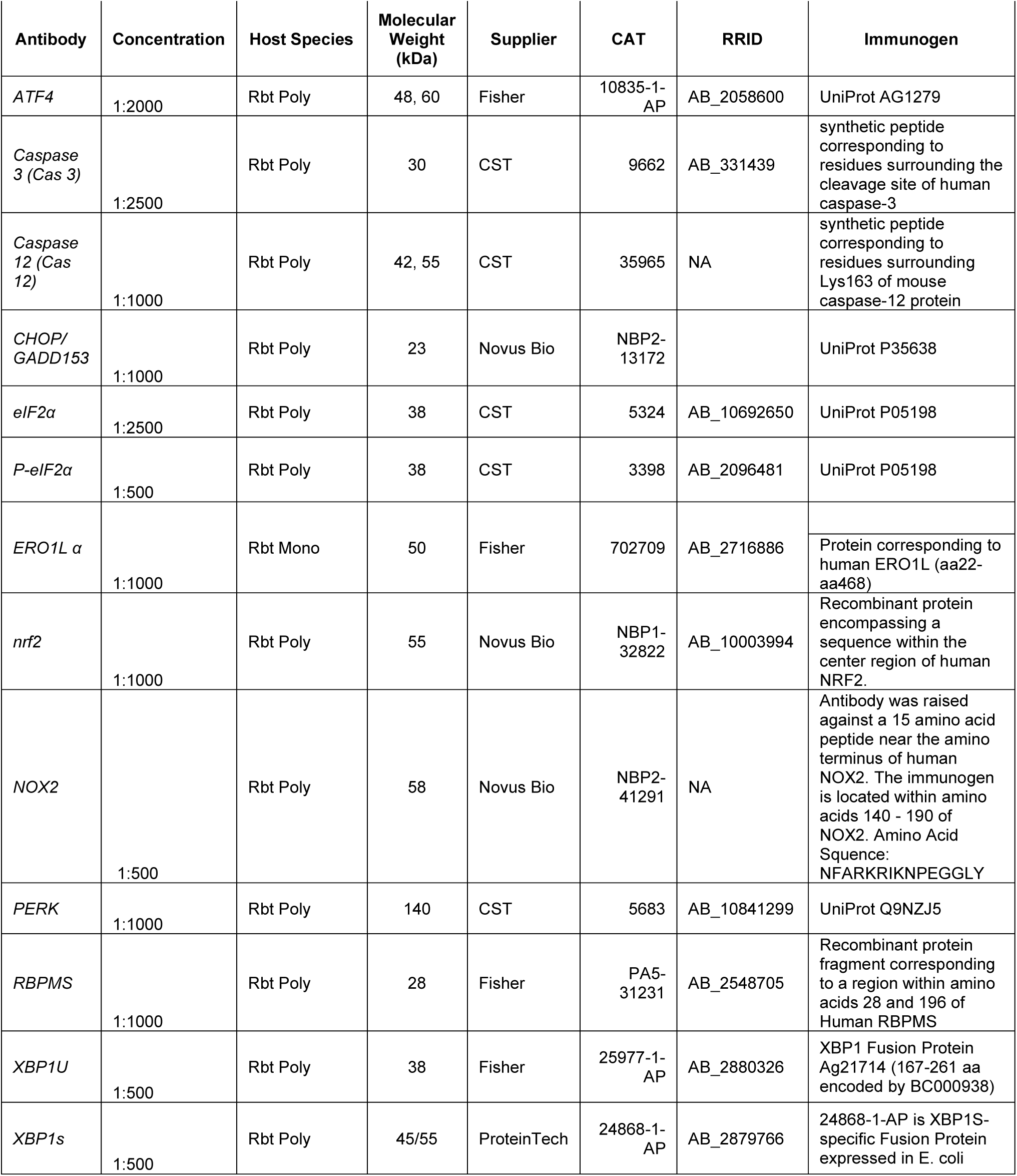
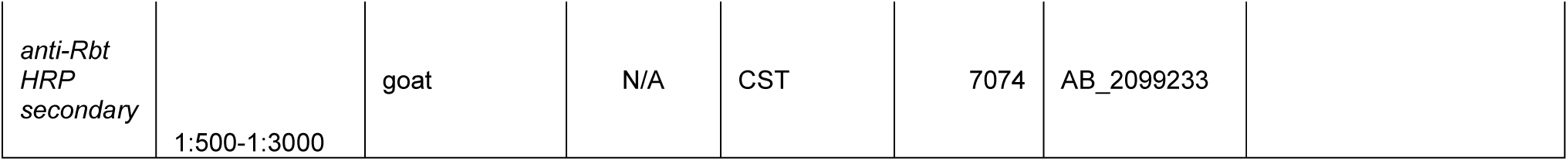
Antibodies.

**Table 3.**
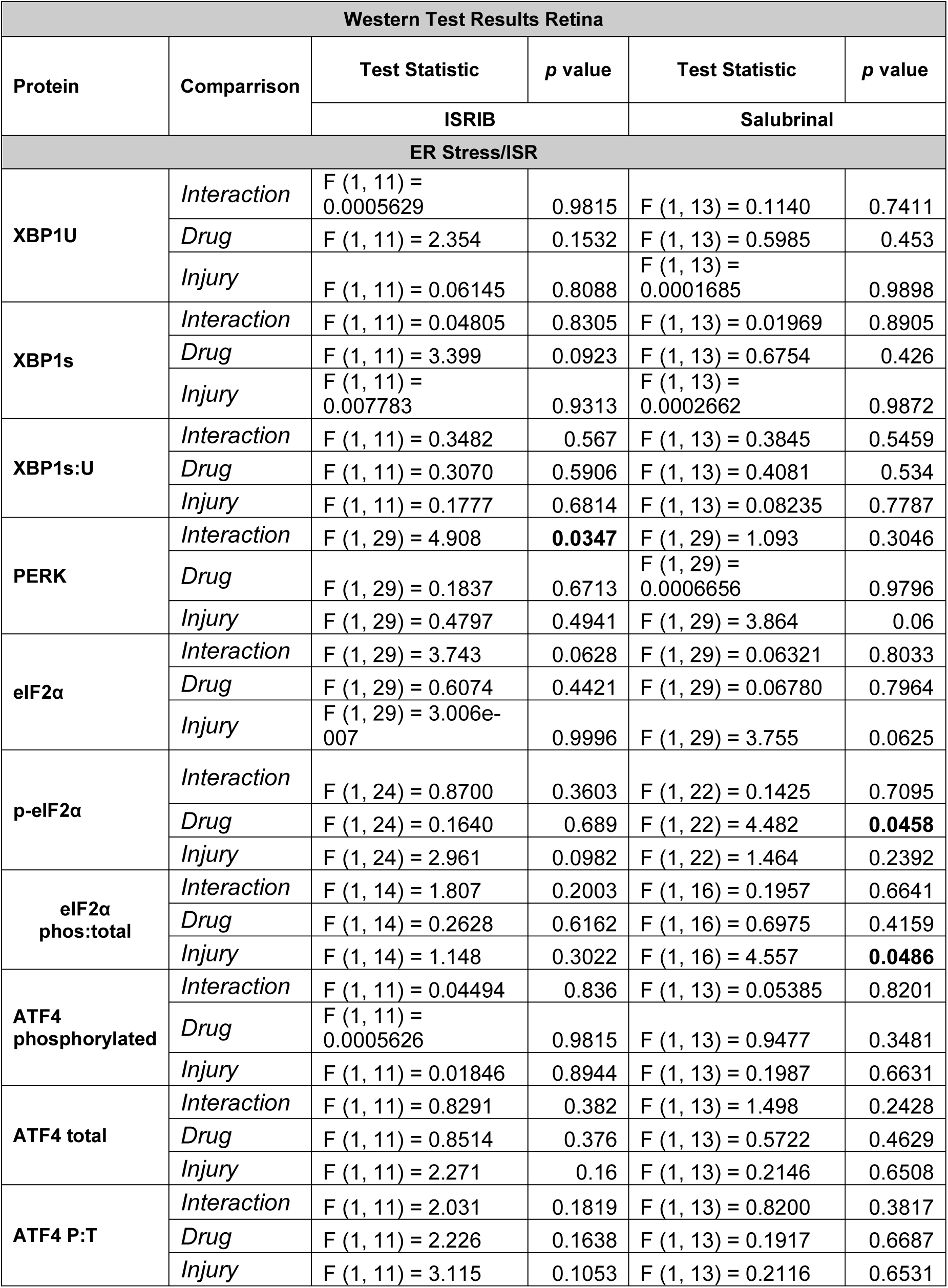
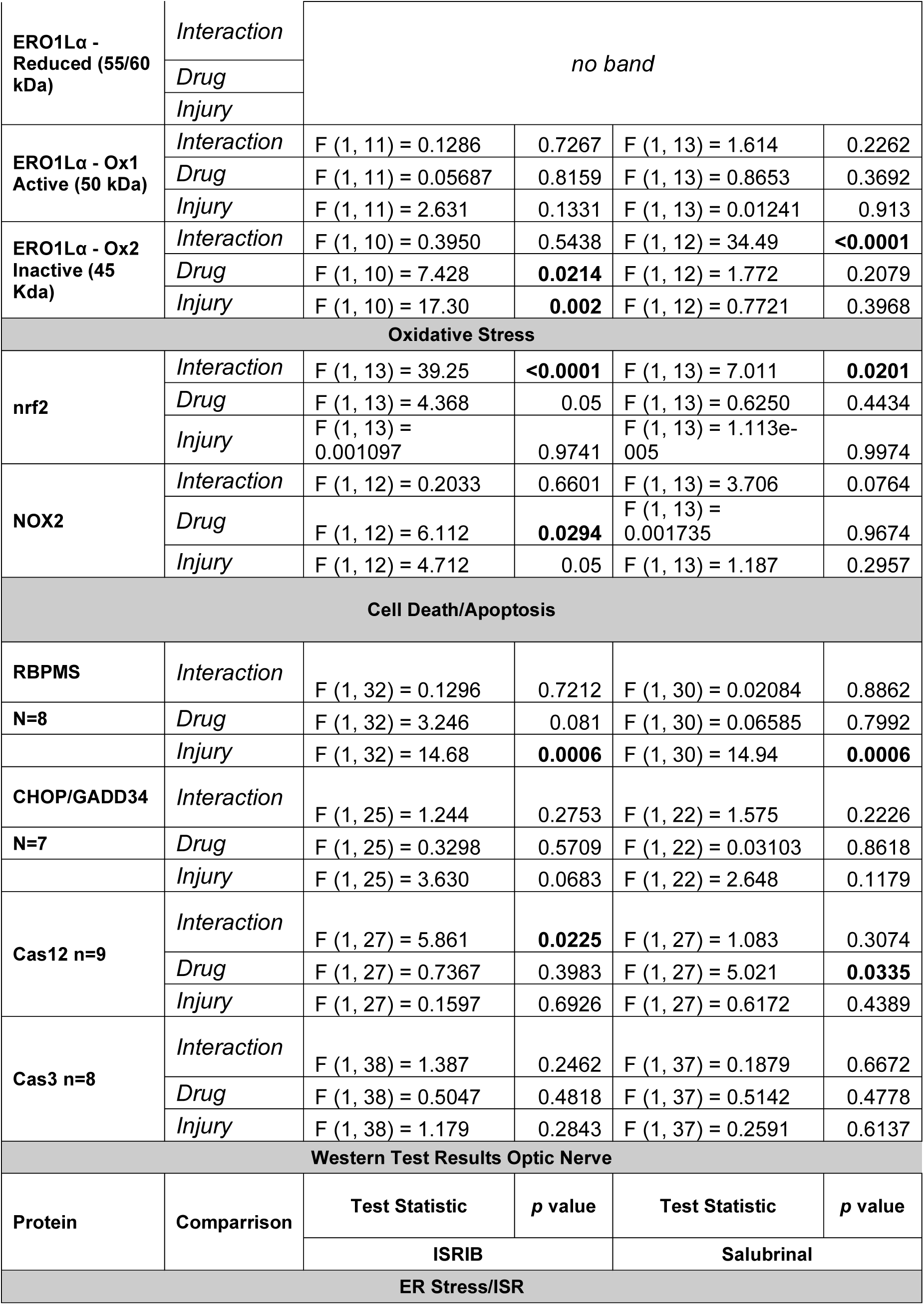
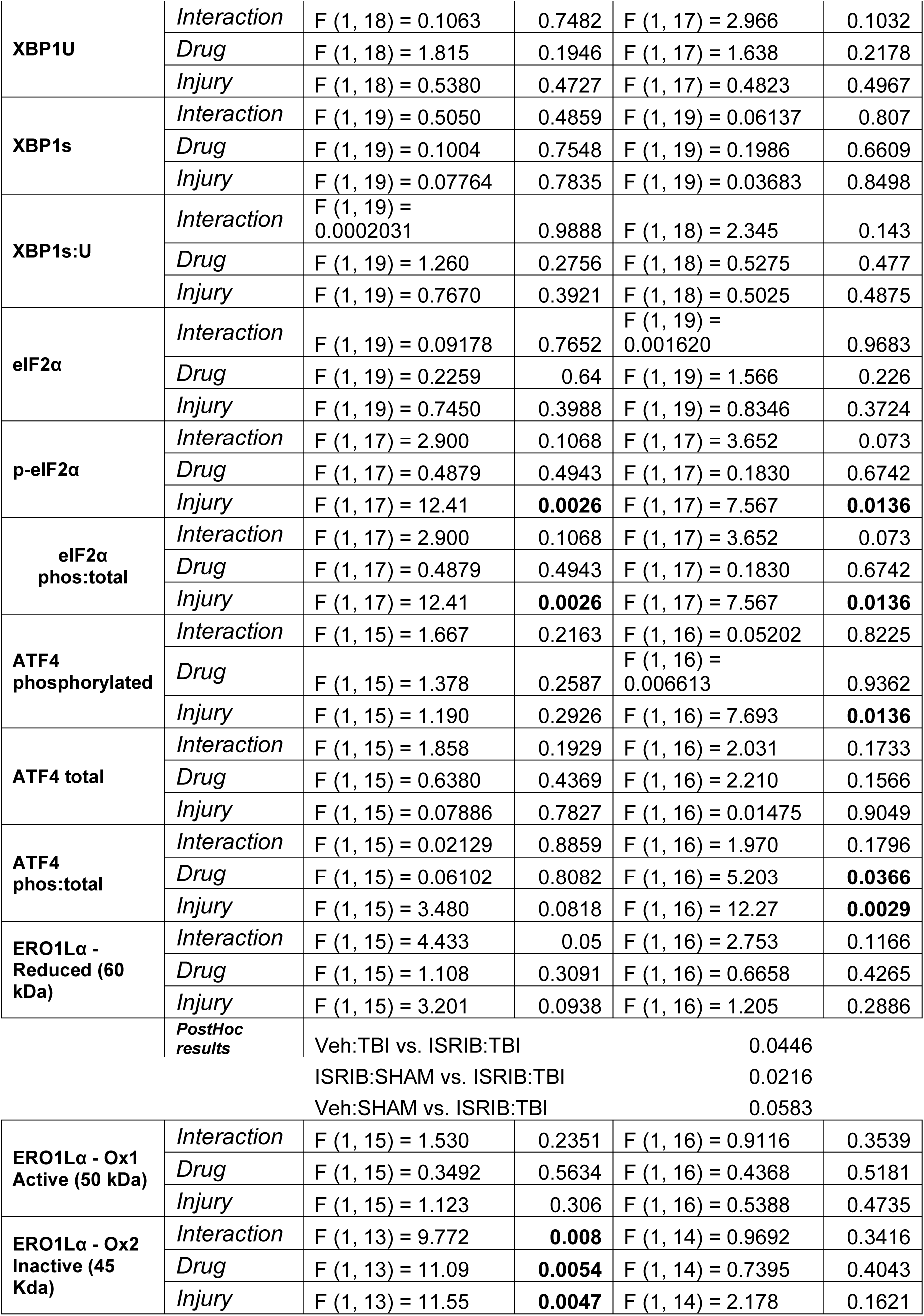
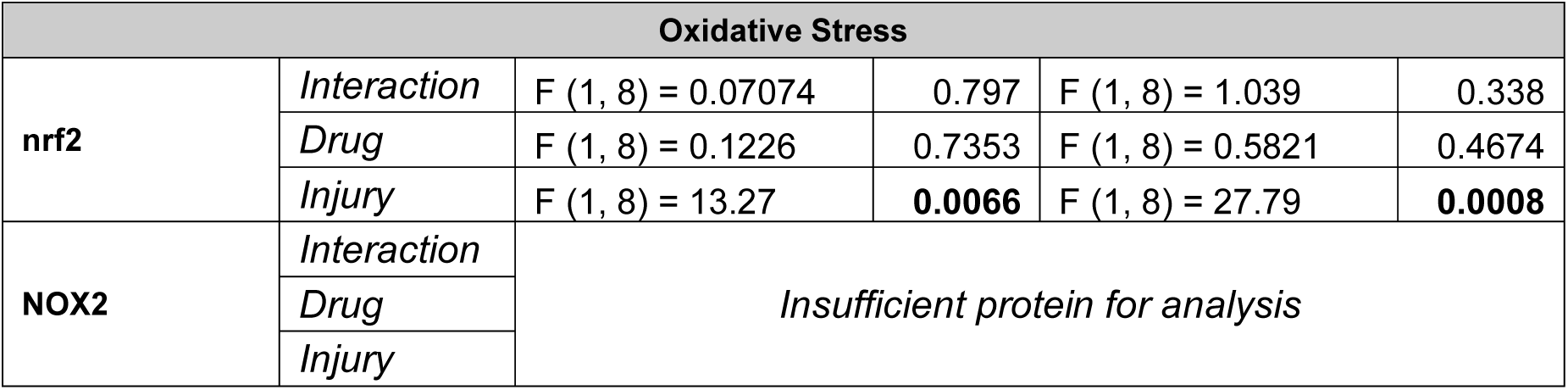
Western Statistics.

### Statistical Analysis

Optomotor behavioral data were analyzed utilizing a 2-way ANOVA within each grating for each drug separately (injury x Sal/Veh or injury x ISRIB/Veh). This is done because responses to individual gratings are expected to differ from each other. Results are presented with each grating in one graph for ease of interpretation. Western blot data were also analyzed by 2-way ANOVA for each protein of interest and graphed with both drugs on the same plot for ease. A critical significance level, α, was set at p≤0.05 for all statistical analyses. Fisher’s LSD was used for post-hoc analyses when relevant.

## Results

### Visual Function after TBI is differentially improved with Salubrinal and ISRIB

We first began our assessment of ER stress inhibition by observing the effects of each drug intervention on overall visual response as indicated by the optomotor behavioral assay. This assay was designed not only to assess overall performance/retention of optomotor nystagmus but also to ascertain any injury-induced drop offs in visual acuity.

Starting with ISRIB intervention (Figure 3a), overall results show no improvement in OMR. However, there is an increase in visual response in TBI+ISRIB mice at higher spatial frequencies (i.e., narrower visual gratings). A main effect of TBI was observed across all gratings (0.12, 0.26, 0.32 and 0.39) with injury significantly reducing the overall number of OMRs recorded. The main effect of drug was found only at 0.26 and 0.32 cpd (“optimal/highest” range of mouse acuity as verified in our previous publications) with a significant interaction at 0.26 and 0.39. The interaction at 0.26 cpd was driven largely by a significant reduction in the total number of OMRs recorded by SHAM mice given ISRIB compared to SHAM+Vehicle (p=0.02) while both injured groups performed significantly worse than both SHAMs. In contrast, the interaction at 0.39 cpd was driven by TBI+ISRIB mice, who were not significantly different from SHAM+ISRIB mice (p=0.6) and performed better than TBI+Vehicle mice (p=0.03), though not significantly. Importantly, SHAM+Vehicle responses were not significantly different from SHAM+ISRIB at all gratings except 0.26 cpd. Table 1 provides a full accounting of all statistics for these data.

Given Salubrinal, similarly non-significant results were found (Figure 3b). A main effect of injury was observed across all gratings of 0.12, 0.26, 0.32 and 0.39. A main effect of drug was only observed at higher spatial resolution gratings (0.32 cpd p=0.02 and 0.39 cpd p=0.04). Like ISRIB, Salubrinal-injected SHAM mice performed worse than vehicle controls at one of the four gratings (0.39) and generally show fewer responses than controls. Despite overall drug effects, there were no improvements between injured vehicle and Salubrinal mice, but there were several instances where injured mice given Salubrinal were no longer impaired compared to SHAM + Salubrinal mice (0.26 and 0.32). Finally, a significant interaction only occurred at 0.39 cpd where Salubrinal induced significant decrease in OMRs between sham groups. Table 1 provides a full accounting of all statistics for these data.

### Salubrinal and ISRIB have long lasting predicted effects on eIF2α phosphorylation in the retina but not in the optic nerve

Injury elevated the ratio of p-eIF2α to total eIF2α in both retinas and optic nerves (Figure 4). There were no significant increases given ISRIB in the retina (p =0.3) nor in the nerve (p=0.07), in line with its mechanism of action. Conversely, Salubrinal treatment is associated with maintained elevated levels of p-eIF2α following TBI, as shown by a main effect of injury (p=0.04) in the retina. However, in the nerve, TBI + Salubrinal mice showed no increase in p-eIF2α ratio despite an increase in TBI + Vehicle mice. Further, there were no outright differences in total eIF2α in either tissue, but in the retina there was a significant increase in p-eIF2α given TBI + Salubrinal (main effect of drug 0.04) (supplementary figure 2a-d). Detailed statistics for all western analyses are presented in table three.

**Figure 4.**
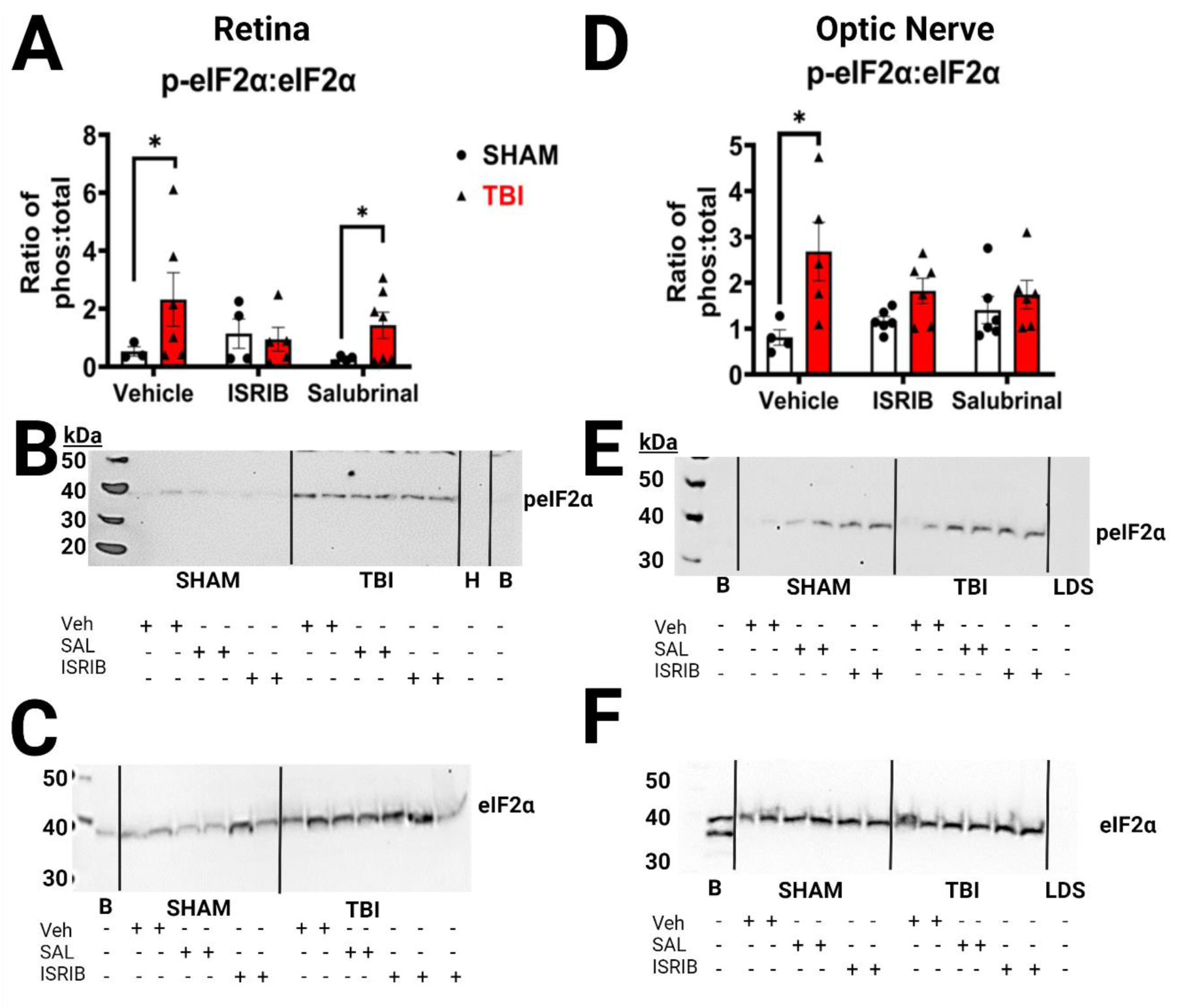
eIF2α phosphorylation in the retina and optic nerve. **(A)** Retinal lysate revealed that both ISRIB and Salubrinal performed as anticipated with long-lasting suppression of eIF2α phosphorylation with ISRIB in injured mice and maintained p-eIF2α with Salubrinal. **(B & C)** Show representative retinal western blots separated by condition with molecular weight (kDa) identification provided using a Magic Mark ladder labeled on the left of the blots. **(D)** Although p-eIF2α was increased with injury, we did not see the same increase in the retina given Salubrinal. **(E & F)** Show representative optic nerve western blots. B=intermembrane control sham brain homogenate, H=H_2_O empty lane, L= ladder, LDS=LDS sample buffer (i.e., negative, empty lane), SAL= Salubrinal, Veh = Vehicle. Data was analyzed by t-tests (injury within drugs) and is presented as mean ± SEM with each symbol representing an individual animal. * p<0.05

### Neither Salubrinal nor ISRIB prevent retinal cell loss or affect apoptosis

RBPMS western blot analysis (figure 5a,b) displayed no retinal ganglion cell preservation given either drug. A main effect of injury was observed as expected except between SHAM and TBI. Although TBI+ISRIB treatment led to a trend toward reduced RBPMS expression compared to SHAM+ISRIB, this was not statistically significant (p=0.06). Additionally, there was no upregulation of apoptotic markers. While our lab has shown upregulation of Caspase-3 at this time point in adolescent mice, this has not been the case with our adult animals (figure 5c, d). However, we do repeatedly find that peak cell death has usually occurred by 7 days post injury with most active cell-death likely complete before this timepoint, so this result is unsurprising. We also measured a well-described pro-apoptotic factor downstream of eIF2α and ATF4, CHOP (C/EBP Homologous Protein; aka GADD153) (Marciniak et al., 2004). We previously reported elevated CHOP expression in this model given injury (Hetzer et al., 2021), and our results support that this may still be true; there is a non-significant trend toward CHOP elevation in TBI+Vehicle compared to for, although not statistically significant in this experiment (p=0.06), TBI+Vehicle CHOP is elevated compared to SHAM (Figure 5 e, f). Because this was not significant, it is difficult to draw firm conclusions, however. Finally, Caspase-12 has been argued to be an ER-specific caspase(Rao et al., 2002), so we looked for elevation in this pro-apoptotic factor. Although there was a significant interaction when assessing ISRIB intervention, post hoc tests revealed no significant t-tests (figure 5g, h). Additionally, a main effect of drug was found for Salubrinal, but post hoc tests revealed no significance. Because these markers are predominantly associated with somatic responses, they were not analyzed in optic nerve samples(Rao et al., 2002; Sanges et al., 2006).

**Figure 5.**
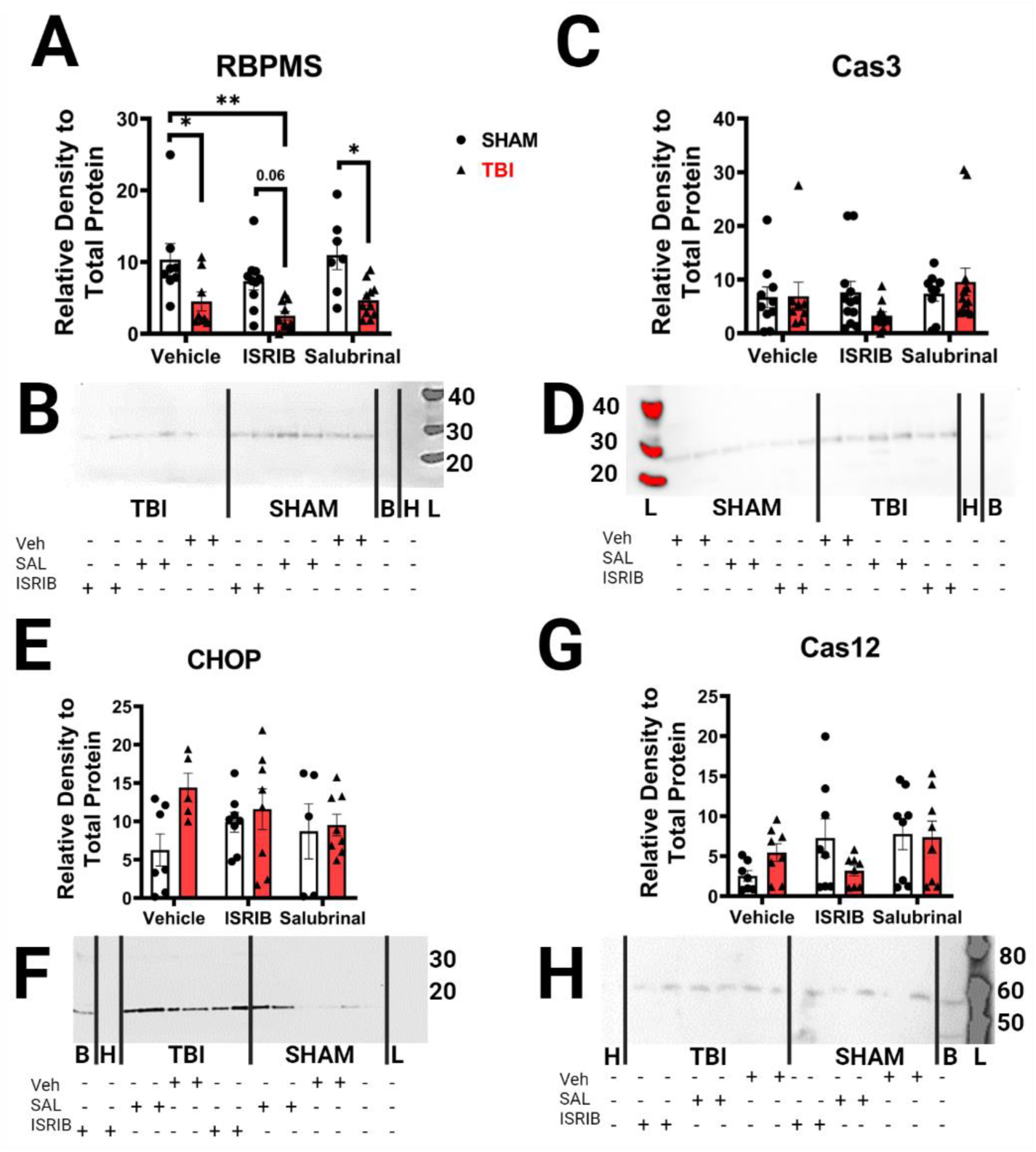
Retinal Apoptosis and RGC protein expression. **(A)** RGC-specific protein RBPMS was decreased in all conditions. Although CHOP expression **(E)** increased with injury, there were no significant differences in any of the apoptosis-related markers examined including **(C)** Caspase 3 **(E)** CHOP or **(G)** Caspase 12. Representative blots for each marker are shown below their respective graphs in **(B,D,F, and H)**. Data was analyzed by t-tests (injury within drugs) and is presented as mean ± SEM with each symbol representing an individual animal. * p<0.05, ** p<0.01

### Only some downstream effectors of the PERK, and not the ER-stress specific IRE1α, pathway are affected by drug, and this is region dependent

Because of the overlapping nature of ER stress with the Integrated Stress response, we wanted to confirm activation of a second ER stress pathway – IRE1α. To do this we examined the downstream X-box Binding protein 1 (XBP1), which is spliced upon IRE1α oligomerization allowing it to upregulate genes involved in both maintenance of proteins and their degradation (Hetz et al., 2020). Additionally, we chose XBP1 as an indicator of ER stress as there is crosstalk and parallel activation of XBP1 by the third branch of the UPR, ATF6.(Walter and Ron, 2011) Unspliced XBP1 (XBP1u) and spliced XBP1 (XBP1s) were not different between any groups in the retina or the nerve (Supplementary Figure 1e-h). We found no difference in the ratio of the two (figure 6a-d).

**Figure 6.**
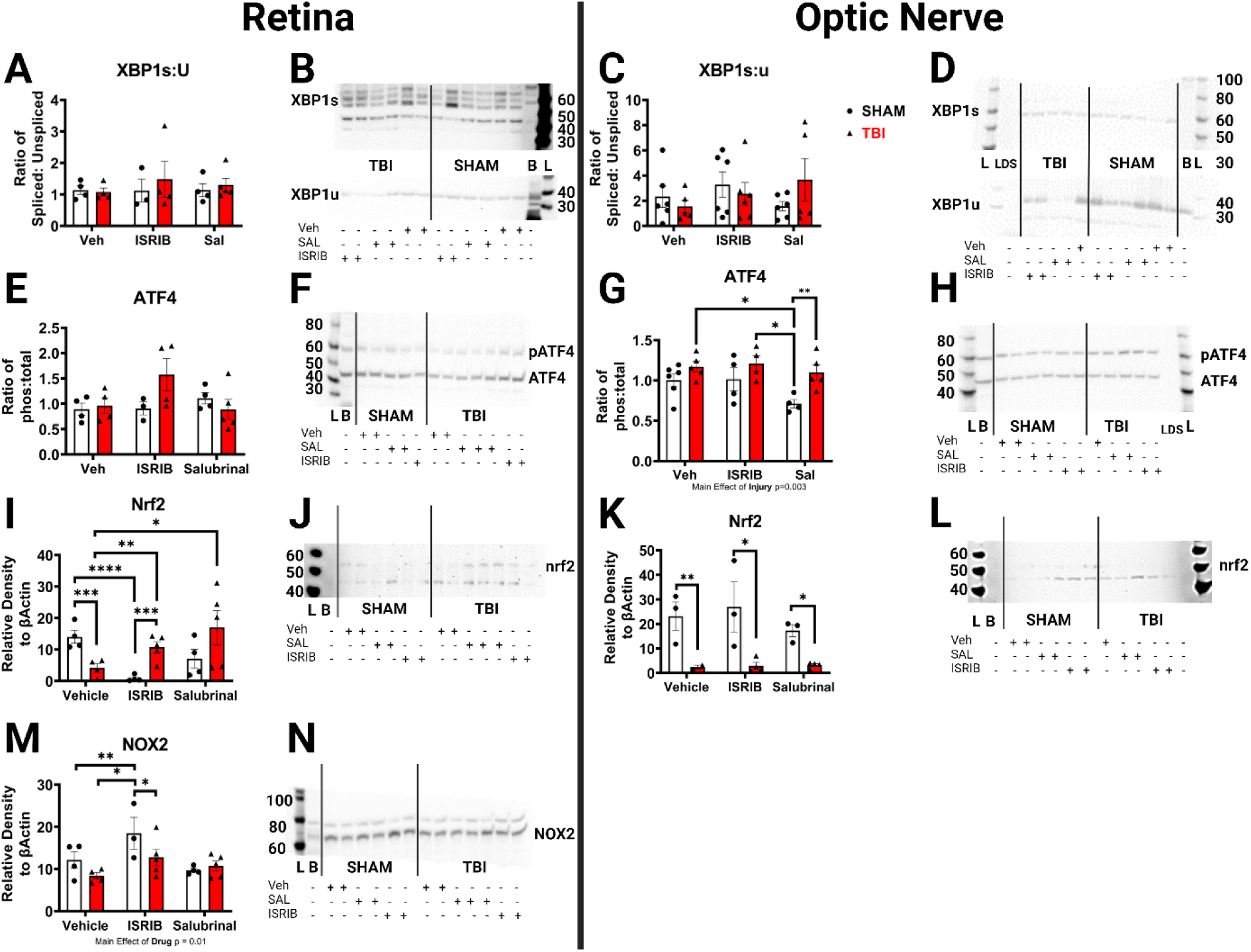
UPR and Oxidative Stress Markers. The ratio of spliced XBP1 to total was not different between groups in retinal samples **(A,B)** or optic nerve samples **(C,D).** The ratio of total to phosphorylated ATF4 was also not changed in the retina **(E,F)** but it was elevated given Salubrinal in the optic nerve **(G,H).** The antioxidant nrf2 was decreased after injury and both ISRIB and Salubrinal increased expression in the retina **(I,J)** but not in the nerve where it remained depressed given injury **(K,L).** ISRIB alone increased the oxidative stress marker NOX2 in control mice and TBI decreased it **(M,N).** There was insufficient optic nerve lysate remaining to assess NOX2 in the optic nerve. B=intermembrane control sham brain homogenate, H=H_2_O empty lane, L= ladder, LDS=LDS empty lane, SAL= Salubrinal, Veh = Vehicle. Data was analyzed by t-tests (injury within drugs) and is presented as mean ± SEM with each symbol representing an individual animal. * p<0.05, ** p<0.01, *** p<0.001, ****p<0.0001.

As previously described, the PERK pathway hinges on eIF2α phosphorylation, after which markers like ATF4 and nrf2 (Nuclear factor erythroid 2-related factor 2) can be upregulated to respond to the stress at hand. ATF4, a transcription factor, can be assessed in total and phosphorylated forms. There were no changes in either form in the retina. Further, this did not change with intervention (Supplementary Figure 1i-j) or as a ratio (Figure 6e-f). In the nerve, we show no change in total ATF4, but a significant main effect of injury (p=0.02) for phosphorylated ATF4 (Supplementary Figure k-l). Moreover, there was a main effect of Salubrinal (p=0.04) and injury (p=0.003) when considering the ratio between the two within-subjects (Figure 6g-h). Post-hoc analyses revealed lower rates of ATF4 phosphorylation given Salubrinal, but only in SHAM mice who had a significantly lower ratio compared to their injured counterparts (p= 0.004). There were no effects of ISRIB on ATF4 expression in the nerve (figure 6g-h).

Further downstream, beneficial factors like nrf2 can be translated to reduce oxidative burden. In the retina, there was a significant interaction for both drugs driven largely by the significant increase in nrf2 expression given both injury and either ISRIB or Salubrinal, though Salubrinal increases nrf2 more drastically (figure 6i-j). A complete inverse was true of the optic nerve, whereby there were main effects of injury revealing a significant reduction in overall nrf2 expression regardless of intervention (figure 6k-l). Because we observed an increase in nrf2 in the retina, we checked for a commonly upregulated oxidative stress marker in TBI – NADPH oxidase 2 (NOX2) (Chuang et al., 2013). There was a main effect of ISRIB (p=0.02), and a main effect of injury (p=0.05). Post-hoc tests revealed a significant increase in NOX2 in SHAM+ISRIB mice compared to TBI + Vehicle (p=0.009) and SHAM + Vehicle (p=0.03), which was then reduced in TBI + ISRIB mice (p=0.04; figure 6m-n). There were no significant differences when using Salubrinal (figure 6m-n). Lack of sufficient sample prevented us from looking at NOX2 in the optic nerve.

### ER protein folding chaperone, ERO1Lα, is differentially affected by injury, drug, and location

Two key protein folding chaperones reside in the ER to produce the necessary disulfide bonds for proper protein folding. We only had sufficient protein to assess one of these chaperones, so we chose ERO1α. If done under non-reducing conditions, three bands can be distinguished via western blot – a low twice-oxidized (Ox2) band at 45 kDa, a middle once-oxidized (Ox1) band around 50 kDa, and a reduced (R) band around 60 kDa (Benham et al., 2013b). In the retina, only the oxidized forms of ERO1α were detected. We show no differences in the Ox1 band for either drug. For ISRIB, we found main effects of both drug (p=0.02) and injury (p=0.002) for the Ox2 band (Figure 7a). Injury significantly reduced overall ERO1α Ox2, but ISRIB was able to significantly increase this between TBI+Veh and TBI+ISRIB (p=0.04). For Salubrinal, there was a significant interaction (p<0.0001). Injury reduced overall ERO1α Ox2 (p=0.0005), and Salubrinal alone decreased ERO1α Ox2 between SHAM vehicle and Salubrinal mice (p=0.007). However, given injury, Salubrinal had the opposite effect whereby it significantly increased ERO1α-Ox2 between TBI + Vehicle and TBI+Salubrinal groups (p=0.0003; Figure 7c).

**Figure 7.**
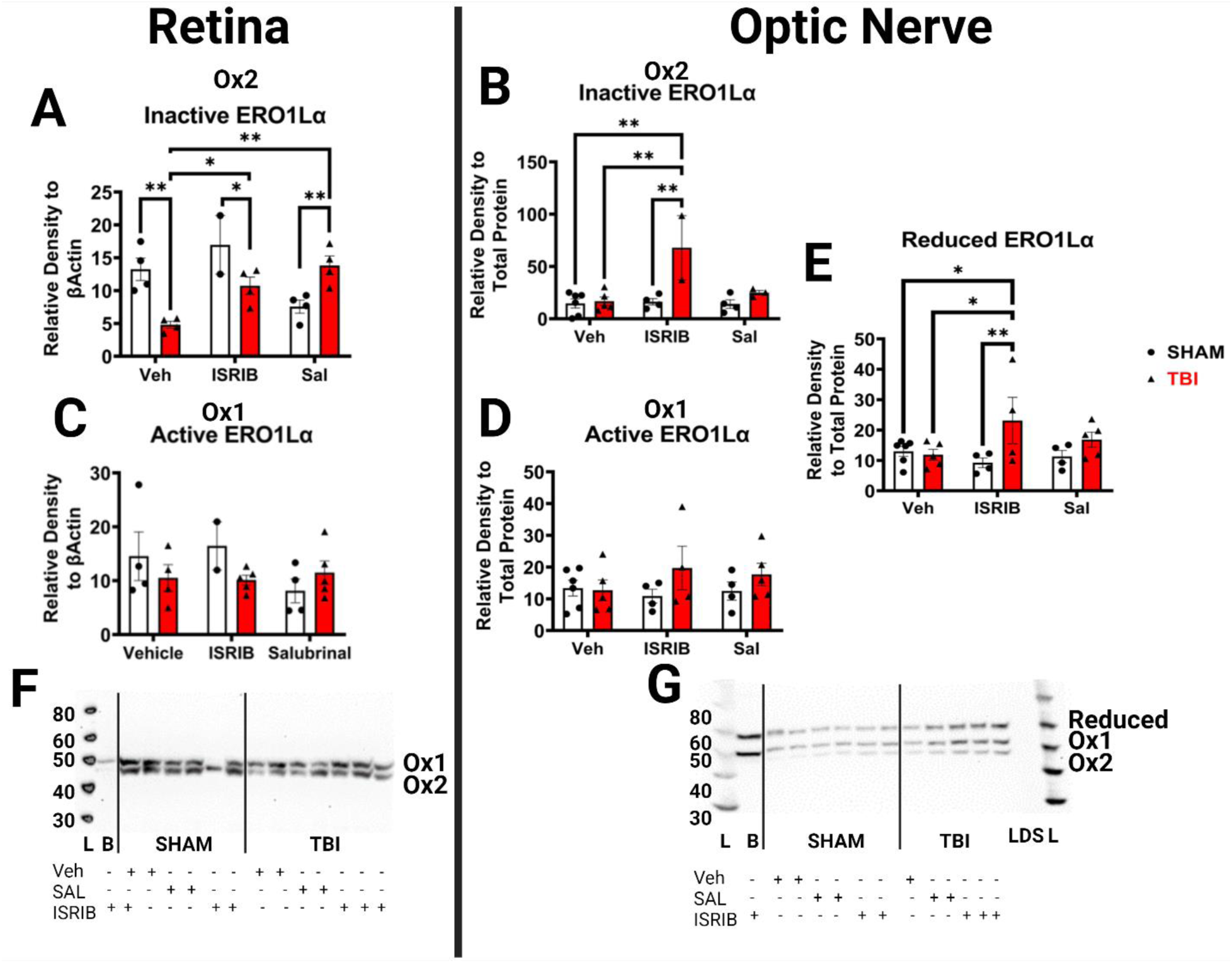
ERO1α expression. In the retina ERO1α-Ox was reduced after injury **(A)** despite no compensatory increase in the active Ox1 variant **(C).** ISRIB increased ERO1α-Ox2 and TBI then decreased it, while Salubrinal increased ERO1α-Ox2 in injured mice **(A)**. In the optic nerve, only ISRIB+TBI increased the Ox2 variant **(B)** and the reduced form or ERO1α **€,** but there were still no changes to the active ERO1α-Ox1 **(D).** Representative blots are provided in for retinal ERO1α **(F)** and optic nerve **(G).** B=intermembrane control sham brain homogenate, L= ladder, LDS=LDS empty lane, SAL= Salubrinal, Veh = Vehicle. Data was analyzed by t-tests (injury within drugs) and is presented as mean ± SEM with each symbol representing an individual animal. * p<0.05, ** p<0.01.

We also found significant changes in ERO1α in the optic nerve. Here, there was a main effect of injury (p=0.01) on ERO1α-Ox2 (Figure 7b). Both injured vehicle (p=0.03) and injured ISRIB (p=0.04) mice had less ERO1α-Ox2 than SHAMs. This reduction was not seen in injured Salubrinal mice (p=0.9). As in the retina there were no changes in the Ox1 band (Figure 7d). However, there was a detectable reduced ERO1α band in optic nerve blots. Analyses revealed no effect of TBI unless given ISRIB, in which case TBI+ISRIB mice had elevated levels of reduced ERO1α compared to TBI+Veh and SHAM+ISRIB (p<0.05) and SHAM + Veh (p=0.05; Figure 7e).

## Discussion

We show that interventions targeting eIF2α in a model of traumatic optic neuropathy do not improve visual function nor retinal ganglion cell survival (at least at the dosages amount and schedule we used) but do reveal intricacies in compartmentalized ER stress responses and effects of ISRIB alone that TBI further impacts. Despite appropriate regulation of eIF2α phosphorylation based on the mechanisms of Salubrinal and ISRIB, we found few significant differences in ER stress response and in downstream integrated stress response factors. Instead, the protein folding machinery of the ER, particularly the primary oxidase ERO1α, appears to be more sensitive to these manipulations. We further show that ER and oxidative stress associated protein expression is unique based on which compartment of the neuron we studied with retinal extracts revealing predicted downstream proteins like nrf2 with upregulation of p-eIF2α while the opposite was true of the optic nerve.

Although some literature suggests that PERK pathway stimulation via interventions like Salubrinal can be neuroprotective via prolonged increases in p-eIF2α (Starr and Gorbatyuk, 2019; Athanasiou et al., 2017) or nrf2 (Comitato et al., 2020), we do not show similar protection in this model. Nor do we show protection when decreasing activation of this pathway, as others have reported with both ISRIB (Chou et al., 2017; Wenzhu Zhou et al., 2023) and other interventions against downstream proteins like ATF4 or CHOP in models of TBI (Hu et al., 2019; Neill and Masson, 2023; Pitale et al., 2017; Song et al., 2008). Optomotor response data showed that ISRIB and Salubrinal led to modest improvements in apparent visual acuity but did not pose a direct benefit on OMRs. These impacts on visual acuity suggest that either drug may play a role in the restoration/preservation of photoreceptors (cones/rods) following injury rather than RGCs. Indeed, ISRIB may preferentially preserve cone function despite loss of rods in a model of inherited retinal degeneration (Gorbatyuk et al., 2020).

Our western blot findings that neither intervention prevented loss of RGCs could also support this line of thinking. Inhibition of the apoptotic factor, CHOP, occurred in both drug conditions despite an increase in TBI+vehicle mice. This indicates that even though these interventions may lower ER-mediated apoptosis at 7 days post injury, this was not sufficient to prevent equal loss of RBPMS expression. This becomes a more likely theory when considering our unpublished electroretinogram findings. In a small cohort of control versus injured mice, we showed significant delays in both A wave (a rod-driven response) and B wave (a bipolar cell/cone driven response) latency at 7 DPI. This deficit was not seen at 24 hours, 3 days, 14 days, or 30 days post injury, suggesting that 7 days post injury is a critical time point for photoreceptors (Supplementary Figure 2). Unfortunately, limited tissue prevented us from determining the effects of our interventions on photoreceptors, which also respond to ER stress and ISRIB treatment (Gorbatyuk et al., 2020).

It is also important to note that there were inhibitory effects of both ISRIB and Salubrinal on SHAM mouse OMR, potentially through undescribed ER stress mechanisms. The combination of injury and ISRIB, in particular, revealed novel interactions. ISRIB treatment alone increased expression of the ER-mediated pro-apoptotic caspase12 and oxidative stress marker, NADPH oxidase 2 (NOX2). Due to the complex nature of the Integrated Stress response, which, similarly to ER stress, hinges on eIF2α phosphorylation, but has many more channels for activation, ISRIB could be dampening the regulatory need of the system for p-eIF2α. This makes sense, because in the eye the environment is more highly oxidative than in other tissues, so it is capable of managing higher levels of oxidative stress. Perhaps inhibiting a pathway for dealing with oxidative stress, protein folding, and metabolic regulation could prove more detrimental in the eye than in the optic nerve. For example, it was found that targeting p-eIF2α, both by systemic GADD34 knockout and photoreceptor specific PERK knock out, in a model of inherited retinal degenerative disease was not protective. Instead, retinal cells might be required to activate alternative pathways for protein synthesis control (Starr and Gorbatyuk, 2019).

Additional studies using models of retinitis pigmentosa in age-related macular degeneration report similar negative findings when p-eIF2α is increased. This contrasts with work performed in hippocampal cultures which showed that Salubrinal-induced upregulation of p-eIF2α and ATF4 led to subsequent antioxidant defense increase and resistance to a number of cellular stressors (Lewerenz and Maher, 2009). Based on our acute findings and these negative effects, it might be that ISRIB/Salubrinal administration may be more harmful to photoreceptors than beneficial for RGCs. Further support for this idea comes from research suggesting that photoreceptors and the retinal pigment epithelium respond poorly to these drugs (McLaughlin et al., 2022) and that basal/homeostatic upregulation of ATF4 can be beneficial for photoreceptor regeneration (Bhootada et al., 2016). This contrasts with the brain or tumor cells where we were unable to find reports of similar effects for either drug. In fact, ISRIB is described as being able to avoid affecting unstressed cells (Sidrauski et al., 2015).

This lack of ER stress upregulation, functional improvement, and RGC preservation in the retina nearly led us to move on from this study. But recent research has taken notice of proximal axon signaling, and it inspired us to probe our collected optic nerve samples. This seemed more important to do once we showed that our model provides a unique opportunity to dissect out proximal axon responses over distal, degenerative mechanisms because axon breakage occurs more distal to the eye than in most other optic nerve injury models. While optic nerve crush, transection, and ocular blast injures optic nerve axons in or behind the eye, we injure the optic nerve in a replicable 1-1.5 mm section just proximal to the optic chiasm.(Hetzer et al., 2023) Thus, most of what we know about the proximal axon is from *in vitro* or *ex vivo* studies (Almasieh et al., 2017; Hao et al., 2016; Hao et al., 2019; Baleriola et al., 2014; Mok et al., 2009; Pathak et al., 2016). This distinction is made more important because differences between signaling in distal and proximal axon segments have been unearthed including retrograde phosphatidyl serine externalization (Almasieh et al., 2017), upregulation of the DLK-cJUN pathway (Asghari Adib et al., 2018; Kievit, 2019; Ugbode et al., 2019; Larhammar et al., 2017), local calcium signaling (Frati et al., 2017; Calixto et al., 2012), and more. It has also been proposed that the autonomous ER may be a critical mechanism for injury signaling in the peripheral nervous system where models of axon injury are made easier to study than in the central nervous system (Farley and Watkins, 2018; Ohtake et al., 2018; Ying et al., 2014). This distinction is important because one process can be regenerative in the PNS, while the same process is degenerative in the CNS (Farley and Watkins, 2018).

Therefore, our ability to study the proximal portion of the optic nerve over the degenerating distal end in the brain, distal to the optic chiasm, could reveal unique ER-stress related injury responses. We might expect some differences between smooth and rough ER as rough ER, predominantly associated with ER stress and protein synthesis (Almanza et al., 2019; Kuijpers et al.; Pryme, 1986), is located only in the soma. In contrast, the axon contains only smooth ER with free floating ribosomes (Kuijpers et al.; Voeltz et al., 2002). Ozturk, et al. (2020) have shown that axonal ER might be less involved in the canonical ER stress response and might, instead, respond to perturbations via Ca^2+^ signaling, lipid synthesis, or communication with mitochondria at mitochondrial-associated membranes (Villegas et al., 2014; Öztürk et al., 2020; Leitman et al., 2013; Carreras-Sureda et al., 2019; Fan and Simmen, 2019; Sun et al., 2017). Furthermore, an argument can be made for the importance of this interconnected meshwork of ER in axonal degeneration as axonal ER can participate in anterograde and retrograde transport and has processes necessary to repair or communicate breakage along the whole of the neuron (Öztürk et al., 2020). It is, therefore, likely that differences exist in how that axon might respond to injury and interventions against ER-stress signaling compared to the soma.

We first assessed eIF2α phosphorylation in the nerve and found increased phosphorylated eIF2α given injury, while ISRIB seemed to reduce this phosphorylation as predicted. However, Salubrinal either had no effect or also reduced p-eIF2α as TBI and sham mice were not different from each other. As in the retina, there was no increase in XBP1s, so we began to look at PERK’s downstream cascade. One of the first factors that eIF2α upregulates upon phosphorylation is ATF4, which targets genes involved in apoptotic, autophagy, antioxidant, and other stress response functions. We previously showed that increased phosphorylation of ATF4, likely indicative of ATF degradation (Lassot et al., 2001), was associated with less axonal degeneration in the optic tract given a brief-oxygen intervention (Torrens et al., 2023). Here we show a similar increase in ATF4 phosphorylation rather than total ATF4 in the optic nerve after Salubrinal treatment. This is interesting because prolonged eIF2α phosphorylation typically leads to prolonged upregulation of total ATF4 translation. However, a recent meta-analysis of ATF4 post translational modifications shows that ATF4 phosphorylation at serine 215, rather than serine 219, increases ATF4 activation/activity (Neill and Masson, 2023). Conversely, we found a trend toward increase in ATF4 phosphorylation in the retina after ISRIB + Injury which makes sense if one follows the previously proposed line of thinking (i.e., reduced p-eIF2α would lead to reduced ATF4 translation or to signals for its degradation upon cessation of ER burden). This difference may have been significant given a higher n or a more sensitive assay. There are luciferase assays for ATF4 that more clearly reveal phosphorylation states and other modifications to better answer the question of what these increases mean that should be considered for future experiments.

This is critical to accurately assess the function of ATF4 in these studies as ATF4’s function is dependent on its binding partners, post-translational modifications, or histone modifications. Within phosphorylation state alone, p-ATF4 could be destined for ubiquitination by SCF E3 ligase (Lassot et al., 2001), decreased transcription of its downstream targets, or overt downregulation (Bagheri-Yarmand et al., 2015; Park et al., 2019). Additionally, we did not collect fixed tissues for imaging in this study, which would be necessary to assess axonal degeneration or changes in the brain. This will be critical in future studies to understand both ER stress localization and whether Salubrinal or ISRIB is more beneficial on the distal side of injury rather than the proximal/somal side that we have analyzed here.

ATF4 can instigate the expression of the antioxidant nrf2, as can the inherent activation of the PERK receptor, tightly linking ER stress to oxidative stress responses. But little is currently known about this ER stress-oxidative stress relationship, particularly under pathological conditions. Studied alone, oxidative stress is a major mechanism of injury pathology associated with TBI (Ismail et al., 2020; Chandran et al., 2021; Khatri et al., 2018; Kontos and Povlishock, 1986), ocular diseases (Masuda et al., 2017; Rohowetz et al., 2018; Chan et al., 2020), and TON (O’Hare Doig et al., 2014; Tao et al., 2017; Bricker-Anthony et al., 2014; Bernardo-Colón et al., 2018; DeJulius et al., 2021). As predicted, Salubrinal most significantly increased nrf2 in injured mice. Perhaps more intriguing is that this increase was not seen in the optic nerve. Although nrf2 requires nuclear translation and this could explain a lack of nrf2 in the axon, it seems odd that it would not be shuttled down the axon to the primary site of injury or expressed in glial cells in the nerve. However, our samples are from seven days post injury, which could be too late to see a need for antioxidants in the nerve. Support for this possibility comes from a variable time course of ROS elevation, which can range from one hour to 28 days (Bricker-Anthony et al., 2014; Lieven et al., 2006; Kanamori et al., 2010; O’Hare Doig et al., 2014), but this research is limited to understanding the retinal response not the axonal response. Future studies will need to probe the axon for ROS markers and perhaps visualize where and when exactly oxidative stress occurs in each compartment of the injured neuron (i.e., somal versus proximal versus distal).

We were able to assess one ROS producing enzyme, Nicotinamide adenine dinucleotide phosphate reduced (NADPH) oxidase 2 (NOX2). NOXs have been associated with TBI and reviewed by many (Khatri et al., 2018; Yao et al., 2021). Compared to other NOXs after TBI, NOX2 appears most consistently upregulated in the central nervous system (Chuang et al., 2013), where it is localized in the plasma membrane of neurons and glial cells (Ibi and Yabe-Nishimura, 2020). Moreover, ER stress mediated NOX2 upregulation is more likely to lead to cell death than other NOXs like NOX4, which can be both pro-survival and proapoptotic (Laurindo et al., 2014). As mentioned above, ISRIB treatment significantly increased NOX2 expression in control mice while TBI nearly significantly decreased overall NOX2 expression with or without ISRIB treatment. Perhaps using ISRIB when there is no increase in ER/oxidative stress prevents the homeostatic functions of this combined system. When ER stress would otherwise have led to oxidative stress, ISRIB might be able to prevent NOX2 expression. Support for linkage of these two pathways comes from a study by Li et al., showing that genetic deletion of NOX2 blocked ER-mediated apoptosis (Li et al., 2010), similar to our finding of blocked CHOP increase after ISRIB treatment. Here we must note that retinal and optic nerve tissue samples were limited, so by the time these questions became relevant, our sample numbers were reduced.

Like all biological pathways, overlap is bound to arise, and roads point to a critical relationship between ER stress and oxidative stress. Not only do NOXs generate hydrogen peroxide but so does the increase in protein folding that occurs as the ER tries to rebalance the unfolded protein load as ER stress ramps up. The ER folding chaperones ERO1α and protein disulfide isomerase (PDI) create H_2_O_2_ upon disulfide bridge formation during folding, and PDI is also sensitive to NOX signaling (Laurindo et al., 2014; Zito, 2015). Thus, it is surprising that we found no upregulation in NOX2 despite changes in ERO1α. ERO1α is critical for the maintenance of the ER’s oxidizing environment (Pollard et al., 1998) where a too-reducing environment can disrupt bond formation and a too-oxidizing environment can induce improper/excess folding (Sevier and Kaiser, 2008). ERO1α exists in three recognized states: an inactive oxidized (Ox2) form, an active partially oxidized (Ox1) form, and a reduced (R) form. When assessed by western blot under non-reducing conditions, each of these isoforms can be identified (Benham et al., 2013a). In response to a changing redox environment, ERO1α must fluctuate between active and inactive states, which it can do within minutes, positioning it as a crucial indicator of the ER’s redox environment (Benham et al., 2013a).

In the retina, where the rough ER resides and the brunt of protein folding takes place, we found that there is no change in the active Ox1 form of ERO1α, nor is there detectible reduced ERO1α. Instead, injury significantly reduced the inactive form of ERO1α and both ISRIB and Salubrinal increased the inactive Ox2 form of ERO1α. It is odd that there is not an accompanying increase in active/reduced ERO1α with this decrease in the inactive isoform given injury. Still, an increase in Ox2 with ISRIB and Salubrinal suggests that these interventions influence the return of ER to equilibrium as higher levels of inactive Ox2 indicate an increasing oxidizing environment. Increased ERO1α -Ox2 might also imply that there is reduced oxidative stress as decreased ability of ERO1α to interact with PDI would lead to reduced production of H2O2 and increased reduced glutathione (GSH). It has also been suggested that when the ER becomes over oxidized by excess reactive oxygen species, ERO1α-Ox2 dissociation from PDI allows ERO1α to travel to the Golgi complex and further reduce the hyper-oxidation of the ER under duress (Kakihana et al., 2013). Future experiments will need to clarify the binding state and localization of ERO1α to better understand its role in the ER stress response.

In the axon, it is theorized that ERO1α may be less crucial to protein folding maintenance and more important for Ca^2+^ maintenance (Anelli et al., 2011; Li et al., 2009) since smooth ER is enriched with mitochondrial ER-associated membranes (MAMs). In these regions, ERO1α and PERK have been shown to interact with each other to regulate ER-mitochondrial calcium dynamics and metabolically adapt mitochondria via increased MAM contact sites under ER stress in an attempt to restore/moderate ROS levels (Bassot et al., 2023). It is, therefore, interesting that we see no changes in axonal ERO1α after injury unless ISRIB is administered. ISRIB decreased phosphorylation of eIF2α, which would prevent eIF2α’s ability to inhibit protein translation, allowing the ER to continue producing and folding proteins under ER stress and reducing PERK activation. We showed that ISRIB prevents p-eIF2α increase in the nerve, so it is possible that the increase in ERO1α-Ox2 and reduced ERO1α, could indicate that the ER is unable to control the redox environment under these conditions. Future studies will need to better assess PERK changes, MAMs, calcium flux, and the redox state of the ER.

## Limitations

Overall, these data helped us to confirm the acute (i.e., 7 day) effects of Salubrinal and ISRIB on eIF2α and on visual system functioning. Moving forward with this study, the first thing that needs to be reexamined is the dose-response and target specificity. This study utilized a single dose that occurred one-hour post-TBI. Salubrinal has a half-life of 1.2 hours in vivo (Zhang et al., 2012) and ISRIB has a half-life of about 8 hours (Sidrauski et al., 2013). Our tissues were extracted 7 days after injury/injection occurred. Thus, multiple dose administration needs to be explored as well as a dose response to assess whether more pronounced effects could be seen including improved RGC and photoreceptor survival/function. Further, Salubrinal and ISRIB were delivered systemically, and western blot analysis was performed in whole tissue lysate. It is possible that other retinal cells and glia play a role in ER stress responses to injury as several studies have reported both photoreceptor (Gorbatyuk et al., 2020) and glial (McMahon et al., 2012; Chen et al., 2016; Logsdon et al., 2016; Huang et al., 2021; Goodall et al., 2010; Martinon et al., 2010) ER stress responses after injury or disease. To elucidate cell-type specific responses to PERK pathway inhibition, more targeted genetic strategies might also be revealing in this model of TON.

Next, it should be noted that while cell death markers such as Caspase 3 and Caspase 12 were examined using western blotting during this report, these caspases would ideally be analyzed using a more accurate enzyme assay. Caspases are proteolytic enzymes, and while western blot analysis is appropriate if completed in conjunction with cleaved to total caspase ratios, an enzyme assay would provide the best analysis of Caspase activation rather than simply the quantity of protein present. Similarly, only RBPMS expression was utilized to measure retinal cells within this study. We were interested in RGCs as they are the cells that undergo axonal damage following TBI, but no other cell types were studied, and our functional analysis cannot necessarily differentiate among cell types. In future studies, electroretinograms would give a more complete picture of retinal cell function.

Further, the use of western blotting as the primary vehicle for ER stress interpretation is a limitation. The semiquantitative nature of western blotting coupled with whole tissue analysis and questions of antibody-specificity (though we did verify all markers with recombinant positive control protein) should be taken into consideration. Future studies would benefit from such additional assessments as visualization of ER damage with electron microscopy that could be further aided with immunodetection of changes to MAMs, PERK, or IP3 receptors. Such data could provide more insight into rough versus smooth ER response to injury since this has not yet been considered in other models of TON. Additional immunohistochemistry could clarify other region-specific expression of ER stress markers.

Finally, it should be noted that this study was performed only in adult male mice. Further studies should examine the effects of TBI-induced TON within a more diverse demographic of animals, including females and adolescents. Both populations could prove informative as we have shown significant differences between adult and adolescent populations in our model. Adolescents appear to produce a more consistent ER stress response after optic nerve injury, while adults, as were used in this study, have less robust upregulation of the IRE1α pathway. This could be a result of differences in protein translation that come with age, whereby adolescents have a higher rate of protein synthesis and turnover than adults which could be more sensitive to axon injury (Kim et al., 2023). Moreover, we have yet to characterize effects of TON in females, but others have found that oxidative stress responses after TBI, particularly involved with mitochondrial dysfunction (Gupte et al., 2019), are affected by sex through inherent antioxidant properties of female hormones, which could influence ER stress and the redox state of the ER (Morrow et al., 1992; Bayir et al., 2004).

## Conclusions

Overall, our data suggested that treatment with either Salubrinal or ISRIB regulates PERK pathway activity long after they have presumably been fully metabolized. Although this did not result in pronounced functional changes or altered RGC survival, we do see novel alterations in redox-sensitive factors that suggest that the primary mechanism of axon signaling in TON is more likely the integrated stress response or a combination of both ER and oxidative stress responses. These data also show that the eye is sensitive to changes in the PERK pathway since interventions in control mice both worsened performance and increased oxidative and apoptotic markers in the absence of injury. We further confirm that axon and soma do not respond identically to axon injury. Of import to past and future treatment strategies, this difference may be critical to understanding why a therapy might work in one scenario but not another. Thus, compartmentalization of neuronal signaling mechanisms will be an important piece of the puzzle as we continue searching for treatments for TON and traumatic axonal injury.

## Competing Interests

None to declare.

## Funding

This research was funded by the Department of Defense, grant W81XWH-21-1-0907 (NKE); the National Institutes of Health (NIH), grant number NS007453 (SMH); a Cincinnati Children’s Hospital Medical Center Procter Scholar Award (NKE); a University of Cincinnati Dean’s Dissertation Fellowship (SMH); and the Albert J. Ryan Foundation (SMH).

## Data Availability Statement

All data are available upon request to the corresponding author.

## Author Contributions

Conceptualization, S.M.H. and N.K.E.; Methodology, S.M.H. and N.K.E.; Validation, R.B, S.M.H., J.N.T, R.G., and N.K.E.; Formal Analysis, R.B, J.N.T, R.G., and S.M.H.; Investigation, S.M.H., R.B, J.N.T, R.G., and N.K.E.; Resources, N.K.E.; Data Curation, S.M.H. and R.B.; Writing – Original Draft Preparation, S.M.H. and R.B.; Writing – Review & Editing, S.M.H., R.B, J.N.T, R.G., and N.K.E.; Visualization, S.M.H.; Supervision, S.M.H. and N.K.E.; Project administration: S.M.H. and N.K.E.; Funding Acquisition, N.K.E

## Acknowledgments

We would like to acknowledge assistance provided by Jordyn Torrens, Macy Urig, and the lab of Dr. James Herman at the University of Cincinnati.

## Conflicts of Interest

The authors declare no conflict of interest.

## Notes

### Competing Interest Statement

The authors have declared no competing interest.

## References

Almanza, A., Carlesso, A., Chintha, C., Creedican, S., Doultsinos, D., Leuzzi, B., Luís, A., Mccarthy, N., Montibeller, L., More, S., Papaioannou, A., Püschel, F., Sassano, M. L., Skoko, J., Agostinis, P., De Belleroche, J., Eriksson, L. A., Fulda, S., Gorman, A. M., Healy, S., Kozlov, A., Muñoz-Pinedo, C., Rehm, M., Chevet, E. & Samali, A. 2019. Endoplasmic reticulum stress signalling - from basic mechanisms to clinical applications. Febs j, 286, 241–278.

Almasieh, M., Catrinescu, M.-M., Binan, L., Costantino, S. & Levin, L. A. 2017. Axonal Degeneration in Retinal Ganglion Cells Is Associated with a Membrane Polarity-Sensitive Redox Process. The Journal of neuroscience: the official journal of the Society for Neuroscience, 37, 3824–3839.

Anelli, T., Bergamelli, L., Margittai, E., Rimessi, A., Fagioli, C., Malgaroli, A., Pinton, P., Ripamonti, M., Rizzuto, R. & Sitia, R. 2011. Ero1α Regulates Ca2+ Fluxes at the Endoplasmic Reticulum–Mitochondria Interface (MAM). Antioxidants & Redox Signaling, 16, 1077–1087.

Asghari Adib, E., Smithson, L. J. & Collins, C. A. 2018. An axonal stress response pathway: degenerative and regenerative signaling by DLK. Current Opinion in Neurobiology, 53, 110–119.

Athanasiou, D., Aguila, M., Bellingham, J., Kanuga, N., Adamson, P. & Cheetham, M. E. 2017. The role of the ER stress-response protein PERK in rhodopsin retinitis pigmentosa. Hum Mol Genet, 26, 4896–4905.

Bagheri-Yarmand, R., Sinha, K. M., Gururaj, A. E., Ahmed, Z., Rizvi, Y. Q., Huang, S. C., Ladbury, J. E., Bogler, O., Williams, M. D., Cote, G. J. & Gagel, R. F. 2015. A novel dual kinase function of the RET proto-oncogene negatively regulates activating transcription factor 4-mediated apoptosis. J Biol Chem, 290, 11749–61.

Baleriola, J., Walker, Chandler A., Jean, Ying Y., Crary, John F., Troy, Carol M., Nagy, Peter L. & Hengst, U. 2014. Axonally Synthesized ATF4 Transmits a Neurodegenerative Signal across Brain Regions. Cell, 158, 1159–1172.

Bassot, A., Chen, J., Takahashi-Yamashiro, K., Yap, M. C., Gibhardt, C. S., Le, G. N. T., Hario, S., Nasu, Y., Moore, J., Gutiérrez, T., Mina, L., Mast, H., Moses, A., Bhat, R., Ballanyi, K., Lemieux, H., Sitia, R., Zito, E., Bogeski, I., Campbell, R. E. & Simmen, T. 2023. The endoplasmic reticulum kinase PERK interacts with the oxidoreductase ERO1 to metabolically adapt mitochondria. Cell Reports, 42, 111899.

Bastakis, G. G., Ktena, N., Karagogeos, D. & Savvaki, M. 2019. Models and treatments for traumatic optic neuropathy and demyelinating optic neuritis. Developmental Neurobiology, 79, 819–836.

Bayir, H., Marion, D. W., Puccio, A. M., Wisniewski, S. R., Janesko, K. L., Clark, R. S. & Kochanek, P. M. 2004. Marked gender effect on lipid peroxidation after severe traumatic brain injury in adult patients. J Neurotrauma, 21, 1–8.

Benham, A. M., Van Lith, M., Sitia, R. & Braakman, I. 2013b. Ero1–PDI interactions, the response to redox flux and the implications for disulfide bond formation in the mammalian endoplasmic reticulum. Philosophical Transactions of the Royal Society B: Biological Sciences, 368, 20110403.

Bernardo-Colón, A., Vest, V., Clark, A., Cooper, M. L., Calkins, D. J., Harrison, F. E. & Rex, T. S. 2018. Antioxidants prevent inflammation and preserve the optic projection and visual function in experimental neurotrauma. Cell Death & Disease, 9, 1097.

Bhootada, Y., Kotla, P., Zolotukhin, S., Gorbatyuk, O., Bebok, Z., Athar, M. & Gorbatyuk, M. 2016. Limited ATF4 Expression in Degenerating Retinas with Ongoing ER Stress Promotes Photoreceptor Survival in a Mouse Model of Autosomal Dominant Retinitis Pigmentosa. PLOS ONE, 11, e0154779.

Bogorad, A. M., Lin, K. Y. & Marintchev, A. 2018. eIF2B Mechanisms of Action and Regulation: A Thermodynamic View. Biochemistry, 57, 1426–1435.

Bond, S., Lopez-Lloreda, C., Gannon, P. J., Akay-Espinoza, C. & Jordan-Sciutto, K. L. 2020. The Integrated Stress Response and Phosphorylated Eukaryotic Initiation Factor 2α in Neurodegeneration. J Neuropathol Exp Neurol, 79, 123–143.

Bricker-Anthony, C., Hines-Beard, J. & Rex, T. S. 2014. Molecular changes and vision loss in a mouse model of closed-globe blast trauma. Invest Ophthalmol Vis Sci, 55, 4853–62.

Calixto, A., Jara, J. S. & Court, F. A. 2012. Diapause formation and downregulation of insulin-like signaling via DAF-16/FOXO delays axonal degeneration and neuronal loss. PLoS Genet, 8, e1003141.

Carreras-Sureda, A., Jaña, F., Urra, H., Durand, S., Mortenson, D. E., Sagredo, A., Bustos, G., Hazari, Y., Ramos-Fernández, E., Sassano, M. L., Pihán, P., Van Vliet, A. R., González-Quiroz, M., Torres, A. K., Tapia-Rojas, C., Kerkhofs, M., Vicente, R., Kaufman, R. J., Inestrosa, N. C., Gonzalez-Billault, C., Wiseman, R. L., Agostinis, P., Bultynck, G., Court, F. A., Kroemer, G., Cárdenas, J. C. & Hetz, C. 2019. Non-canonical function of IRE1α determines mitochondria-associated endoplasmic reticulum composition to control calcium transfer and bioenergetics. Nature Cell Biology, 21, 755–767.

Chan, J. W., Hills, N. K., Bakall, B. & Fernandez, B. 2019. Indirect Traumatic Optic Neuropathy in Mild Chronic Traumatic Brain Injury. Investigative Ophthalmology & Visual Science, 60, 2005–2011.

Chan, T. C. W., Wilkinson Berka, J. L., Deliyanti, D., Hunter, D., Fung, A., Liew, G. & White, A. 2020. The role of reactive oxygen species in the pathogenesis and treatment of retinal diseases. Experimental Eye Research, 201, 108255.

Chandran, R., Mehta, S. L. & Vemuganti, R. 2021. Antioxidant Combo Therapy Protects White Matter After Traumatic Brain Injury. NeuroMolecular Medicine.

Chen, B., Zhang, H., Zhai, Q., Li, H., Wang, C. & Wang, Y. 2022. Traumatic optic neuropathy: a review of current studies. Neurosurgical Review, 45, 1895–1913.

Chen, Y.-J., Liang, C.-M., Tai, M.-C., Chang, Y.-H., Lin, T.-Y., Chung, C.-H., Lin, F.-H., Tsao, C.-H. & Chien, W.-C. 2017. Longitudinal relationship between traumatic brain injury and the risk of incident optic neuropathy: A 10-year follow-up nationally representative Taiwan survey. Oncotarget, 8, 86924–86933.

Chen, Y., Holstein, D. M., Aime, S., Bollo, M. & Lechleiter, J. D. 2016. Calcineurin β protects brain after injury by activating the unfolded protein response. Neurobiology of Disease, 94, 139–156.

Chou, A., Krukowski, K., Jopson, T., Zhu, P. J., Costa-Mattioli, M., Walter, P. & Rosi, S. 2017. Inhibition of the integrated stress response reverses cognitive deficits after traumatic brain injury. Proc Natl Acad Sci U S A, 114, E6420–e6426.

Chuang, D. Y., Chan, M.-H., Zong, Y., Sheng, W., He, Y., Jiang, J. H., Simonyi, A., Gu, Z., Fritsche, K. L., Cui, J., Lee, J. C., Folk, W. R., Lubahn, D. B., Sun, A. Y. & Sun, G. Y. 2013. Magnolia polyphenols attenuate oxidative and inflammatory responses in neurons and microglial cells. Journal of Neuroinflammation, 10, 786.

Comitato, A., Schiroli, D., Montanari, M. & Marigo, V. 2020. Calpain Activation Is the Major Cause of Cell Death in Photoreceptors Expressing a Rhodopsin Misfolding Mutation. Mol Neurobiol, 57, 589–599.

Dejulius, C. R., Bernardo-Colón, A., Naguib, S., Backstrom, J. R., Kavanaugh, T., Gupta, M. K., Duvall, C. L. & Rex, T. S. 2021. Microsphere antioxidant and sustained erythropoietin-R76E release functions cooperate to reduce traumatic optic neuropathy. Journal of Controlled Release, 329, 762–773.

Evans, L. P., Roghair, A. M., Gilkes, N. J. & Bassuk, A. G. 2021. Visual Outcomes in Experimental Rodent Models of Blast-Mediated Traumatic Brain Injury. Frontiers in Molecular Neuroscience, 14.

Evanson, N. K., Guilhaume-Correa, F., Herman, J. P. & Goodman, M. D. 2018. Optic tract injury after closed head traumatic brain injury in mice: A model of indirect traumatic optic neuropathy. PLOS ONE, 13, e0197346.

Fan, Y. & Simmen, T. 2019. Mechanistic Connections between Endoplasmic Reticulum (ER) Redox Control and Mitochondrial Metabolism. Cells, 8, 1071.

Farley, M. M. & Watkins, T. A. 2018. Intrinsic Neuronal Stress Response Pathways in Injury and Disease. Annual Review of Pathology: Mechanisms of Disease, 13, 93–116.

Frati, A., Cerretani, D., Fiaschi, A. I., Frati, P., Gatto, V., La Russa, R., Pesce, A., Pinchi, E., Santurro, A., Fraschetti, F. & Fineschi, V. 2017. Diffuse Axonal Injury and Oxidative Stress: A Comprehensive Review. International Journal of Molecular Sciences, 18, 2600.

Goodall, J. C., Wu, C., Zhang, Y., Mcneill, L., Ellis, L., Saudek, V. & Gaston, J. S. H. 2010. Endoplasmic reticulum stress-induced transcription factor, CHOP, is crucial for dendritic cell IL-23 expression. Proceedings of the National Academy of Sciences, 107, 17698-17703.

Gorbatyuk, M. S., Starr, C. R. & Gorbatyuk, O. S. 2020. Endoplasmic reticulum stress: New insights into the pathogenesis and treatment of retinal degenerative diseases. Prog Retin Eye Res, 79, 100860.

Gupte, R., Brooks, W., Vukas, R., Pierce, J. & Harris, J. 2019. Sex Differences in Traumatic Brain Injury: What We Know and What We Should Know. Journal of neurotrauma, 36, 3063–3091.

Hao, Y., Frey, E., Yoon, C., Wong, H., Nestorovski, D., Holzman, L. B., Giger, R. J., Diantonio, A. & Collins, C. 2016. An evolutionarily conserved mechanism for cAMP elicited axonal regeneration involves direct activation of the dual leucine zipper kinase DLK. eLife, 5, e14048.

Hao, Y., Waller, T. J., Nye, D. M., Li, J., Zhang, Y., Hume, R. I., Rolls, M. M. & Collins, C. A. 2019. Degeneration of Injured Axons and Dendrites Requires Restraint of a Protective JNK Signaling Pathway by the Transmembrane Protein Raw. The Journal of Neuroscience, 39, 8457–8470.

Hetz, C., Zhang, K. & Kaufman, R. J. 2020. Mechanisms, regulation and functions of the unfolded protein response. Nature Reviews Molecular Cell Biology, 21, 421–438.

Hetzer, S. M., Guilhaume-Correa, F., Day, D., Bedolla, A. & Evanson, N. K. 2021. Traumatic Optic Neuropathy Is Associated with Visual Impairment, Neurodegeneration, and Endoplasmic Reticulum Stress in Adolescent Mice. Cells, 10, 996.

Hetzer, S. M., O’connell, C., Lallo, V., Robson, M. & Evanson, N. K. 2023. Model matters: Differential outcomes in traumatic optic neuropathy pathophysiology between blunt and blast-wave mediated head injuries. bioRxiv, 2023.05.25.542261.

Hu, H., Tian, M., Ding, C. & Yu, S. 2019. The C/EBP Homologous Protein (CHOP) Transcription Factor Functions in Endoplasmic Reticulum Stress-Induced Apoptosis and Microbial Infection. Frontiers in Immunology, 9.

Huang, T.-C., Luo, L., Jiang, S.-H., Chen, C., He, H.-Y., Liang, C.-F., Li, W.-S., Wang, H., Zhu, L., Wang, K. & Guo, Y. 2021. Targeting integrated stress response regulates microglial M1/M2 polarization and attenuates neuroinflammation following surgical brain injury in rat. Cellular Signalling, 85, 110048.

Ibi, M. & Yabe-Nishimura, C. 2020. Chapter 1 - The role of reactive oxygen species in the pathogenic pathways of depression. In: Martin, C. R. & Preedy, V. R. (eds.) Oxidative Stress and Dietary Antioxidants in Neurological Diseases. Academic Press.

Ismail, H., Shakkour, Z., Tabet, M., Abdelhady, S., Kobaisi, A., Abedi, R., Nasrallah, L., Pintus, G., Al-Dhaheri, Y., Mondello, S., El-Khoury, R., Eid, A. H., Kobeissy, F. & Salameh, J. 2020. Traumatic Brain Injury: Oxidative Stress and Novel Anti-Oxidants Such as Mitoquinone and Edaravone. Antioxidants, 9, 943.

Kakihana, T., Araki, K., Vavassori, S., Iemura, S., Cortini, M., Fagioli, C., Natsume, T., Sitia, R. & Nagata, K. 2013. Dynamic regulation of Ero1α and peroxiredoxin 4 localization in the secretory pathway. J Biol Chem, 288, 29586–94.

Kanamori, A., Catrinescu, M.-M., Kanamori, N., Mears, K. A., Beaubien, R. & Levin, L. A. 2010. Superoxide is an associated signal for apoptosis in axonal injury. Brain, 133, 2612–2625.

Karimi, S., Arabi, A., Ansari, I., Shahraki, T. & Safi, S. 2021. A Systematic Literature Review on Traumatic Optic Neuropathy. Journal of Ophthalmology, 2021, 5553885.

Khatri, N., Thakur, M., Pareek, V., Kumar, S., Sharma, S. & Datusalia, A. K. 2018. Oxidative Stress: Major Threat in Traumatic Brain Injury. CNS & Neurological Disorders - Drug Targets-CNS & Neurological Disorders), 17, 689–695.

Kievit, B. J. 2019. Reactive oxygen species and transient receptor potential cation channel vanilloid 1 (TRPV1) as mediators of Wallerian degeneration. Text.

Kim, H. S., Parker, D. J., Hardiman, M. M., Munkácsy, E., Jiang, N., Rogers, A. N., Bai, Y., Brent, C., Mobley, J. A., Austad, S. N. & Pickering, A. M. 2023. Early-adulthood spike in protein translation drives aging via juvenile hormone/germline signaling. Nature Communications, 14, 5021.

Konieczny, A. & Safer, B. 1983. Purification of the eukaryotic initiation factor 2-eukaryotic initiation factor 2B complex and characterization of its guanine nucleotide exchange activity during protein synthesis initiation. J Biol Chem, 258, 3402–8.

Kontos, H. A. & Povlishock, J. T. 1986. Oxygen radicals in brain injury. Cent Nerv Syst Trauma, 3, 257–63.

Kuijpers, M., Nguyen, P. T. & Haucke, V. The Endoplasmic Reticulum and Its Contacts: Emerging Roles in Axon Development, Neurotransmission, and Degeneration. The Neuroscientist, 0, 10738584231162810.

Ladak, A. A., Enam, S. A. & Ibrahim, M. T. 2019. A Review of the Molecular Mechanisms of Traumatic Brain Injury. World Neurosurgery, 131, 126–132.

Larhammar, M., Huntwork-Rodriguez, S., Jiang, Z., Solanoy, H., Sengupta Ghosh, A., Wang, B., Kaminker, J. S., Huang, K., Eastham-Anderson, J., Siu, M., Modrusan, Z., Farley, M. M., Tessier-Lavigne, M., Lewcock, J. W. & Watkins, T. A. 2017. Dual leucine zipper kinase-dependent PERK activation contributes to neuronal degeneration following insult. eLife, 6, e20725.

Lassot, I., Ségéral, E., Berlioz-Torrent, C., Durand, H., Groussin, L., Hai, T., Benarous, R. & Margottin-Goguet, F. 2001. ATF4 degradation relies on a phosphorylation-dependent interaction with the SCF(betaTrCP) ubiquitin ligase. Mol Cell Biol, 21, 2192–202.

Laurindo, F. R. M., Araujo, T. L. S. & Abrahão, T. B. 2014. Nox NADPH Oxidases and the Endoplasmic Reticulum. Antioxidants & Redox Signaling, 20, 2755–2775.

Leitman, J., Ron, E., Ogen-Shtern, N. & Lederkremer, G. Z. 2013. Compartmentalization of endoplasmic reticulum quality control and ER-associated degradation factors. DNA Cell Biol, 32, 2–7.

Lewerenz, J. & Maher, P. 2009. Basal levels of eIF2alpha phosphorylation determine cellular antioxidant status by regulating ATF4 and xCT expression. The Journal of biological chemistry, 284, 1106–1115.

Li, G., Mongillo, M., Chin, K.-T., Harding, H., Ron, D., Marks, A. R. & Tabas, I. 2009. Role of ERO1-α–mediated stimulation of inositol 1,4,5-triphosphate receptor activity in endoplasmic reticulum stress–induced apoptosis. Journal of Cell Biology, 186, 783-792.

Li, G., Scull, C., Ozcan, L. & Tabas, I. 2010. NADPH oxidase links endoplasmic reticulum stress, oxidative stress, and PKR activation to induce apoptosis. Journal of Cell Biology, 191, 1113–1125.

Lieven, C. J., Hoegger, M. J., Schlieve, C. R. & Levin, L. A. 2006. Retinal Ganglion Cell Axotomy Induces an Increase in Intracellular Superoxide Anion. Investigative Ophthalmology & Visual Science, 47, 1477–1485.

Logsdon, A. F., Lucke-Wold, B. P., Nguyen, L., Matsumoto, R. R., Turner, R. C., Rosen, C. L. & Huber, J. D. 2016. Salubrinal reduces oxidative stress, neuroinflammation and impulsive-like behavior in a rodent model of traumatic brain injury. Brain Res, 1643, 140–51.

Marciniak, S. J., Yun, C. Y., Oyadomari, S., Novoa, I., Zhang, Y., Jungreis, R., Nagata, K., Harding, H. P. & Ron, D. 2004. CHOP induces death by promoting protein synthesis and oxidation in the stressed endoplasmic reticulum. Genes Dev, 18, 3066–77.

Martinon, F., Chen, X., Lee, A. H. & Glimcher, L. H. 2010. TLR activation of the transcription factor XBP1 regulates innate immune responses in macrophages. Nat Immunol, 11, 411–8.

Masuda, T., Shimazawa, M. & Hara, H. 2017. Retinal Diseases Associated with Oxidative Stress and the Effects of a Free Radical Scavenger (Edaravone). Oxidative Medicine and Cellular Longevity, 2017, 9208489.

Mclaughlin, T., Medina, A., Perkins, J., Yera, M., Wang, J. J. & Zhang, S. X. 2022. Cellular stress signaling and the unfolded protein response in retinal degeneration: mechanisms and therapeutic implications. Mol Neurodegener, 17, 25.

Mcmahon, J. M., Mcquaid, S., Reynolds, R. & Fitzgerald, U. F. 2012. Increased expression of ER stress- and hypoxia-associated molecules in grey matter lesions in multiple sclerosis. Mult Scler, 18, 1437–47.

Mok, S. A., Lund, K. & Campenot, R. B. 2009. A retrograde apoptotic signal originating in NGF-deprived distal axons of rat sympathetic neurons in compartmented cultures. Cell Res, 19, 546–60.

Morrow, J. D., Awad, J. A., Boss, H. J., Blair, I. A. & Roberts, L. J., 2nd 1992. Non-cyclooxygenase-derived prostanoids (F2-isoprostanes) are formed in situ on phospholipids. Proc Natl Acad Sci U S A, 89, 10721-5.

Neill, G. & Masson, G. R. 2023. A stay of execution: ATF4 regulation and potential outcomes for the integrated stress response. Frontiers in Molecular Neuroscience, 16.

O’Hare Doig, R. L., Bartlett, C. A., Maghzal, G. J., Lam, M., Archer, M., Stocker, R. & Fitzgerald, M. 2014. Reactive species and oxidative stress in optic nerve vulnerable to secondary degeneration. Exp Neurol, 261, 136–46.

Ohtake, Y., Matsuhisa, K., Kaneko, M., Kanemoto, S., Asada, R., Imaizumi, K. & Saito, A. 2018. Axonal Activation of the Unfolded Protein Response Promotes Axonal Regeneration Following Peripheral Nerve Injury. Neuroscience, 375, 34–48.

Öztürk, Z., O’kane, C. J. & Pérez-Moreno, J. J. 2020. Axonal Endoplasmic Reticulum Dynamics and Its Roles in Neurodegeneration. Front Neurosci, 14, 48.

Park, D., Gu, H., Baek, J. H. & Baek, K. 2019. Undercarboxylated osteocalcin downregulates pancreatic lipase expression in an ATF4-dependent manner in pancreatic acinar cells. Bone, 127, 220–227.

Pathak, G. K., Ornstein, H., Aranda-Espinoza, H., Karlsson, A. J. & Shah, S. B. 2016. Increases in Retrograde Injury Signaling Complex-Related Transcripts in Central Axons following Injury. Neural Plast, 2016, 3572506.

Pitale, P. M., Gorbatyuk, O. & Gorbatyuk, M. 2017. Neurodegeneration: Keeping ATF4 on a Tight Leash. Frontiers in Cellular Neuroscience, 11.

Pollard, M. G., Travers, K. J. & Weissman, J. S. 1998. Ero1p: A novel and ubiquitous protein with an essential role in oxidative protein folding in the endoplasmic reticulum. Molecular Cell, 1, 171–182.

Pryme, I. F. 1986. Compartmentation of the rough endoplasmic reticulum. Mol Cell Biochem, 71, 3–18.

Rao, R. V., Peel, A., Logvinova, A., Del Rio, G., Hermel, E., Yokota, T., Goldsmith, P. C., Ellerby, L. M., Ellerby, H. M. & Bredesen, D. E. 2002. Coupling endoplasmic reticulum stress to the cell death program: role of the ER chaperone GRP78. FEBS Lett, 514, 122–8.

Rohowetz, L. J., Kraus, J. G. & Koulen, P. 2018. Reactive Oxygen Species-Mediated Damage of Retinal Neurons: Drug Development Targets for Therapies of Chronic Neurodegeneration of the Retina. International Journal of Molecular Sciences, 19, 3362.

Rubovitch, V., Barak, S., Rachmany, L., Goldstein, R. B., Zilberstein, Y. & Pick, C. G. 2015. The neuroprotective effect of salubrinal in a mouse model of traumatic brain injury. Neuromolecular Med, 17, 58–70.

Sanges, D., Comitato, A., Tammaro, R. & Marigo, V. 2006. Apoptosis in retinal degeneration involves cross-talk between apoptosis-inducing factor (AIF) and caspase-12 and is blocked by calpain inhibitors. Proc Natl Acad Sci U S A, 103, 17366–71.

Sevier, C. S. & Kaiser, C. A. 2008. Ero1 and redox homeostasis in the endoplasmic reticulum. Biochimica et Biophysica Acta (BBA) - Molecular Cell Research, 1783, 549–556.

Sidrauski, C., Acosta-Alvear, D., Khoutorsky, A., Vedantham, P., Hearn, B. R., Li, H., Gamache, K., Gallagher, C. M., Ang, K. K., Wilson, C., Okreglak, V., Ashkenazi, A., Hann, B., Nader, K., Arkin, M. R., Renslo, A. R., Sonenberg, N. & Walter, P. 2013. Pharmacological brake-release of mRNA translation enhances cognitive memory. Elife, 2, e00498.

Sidrauski, C., Mcgeachy, A. M., Ingolia, N. T. & Walter, P. 2015. The small molecule ISRIB reverses the effects of eIF2α phosphorylation on translation and stress granule assembly. eLife, 4, e05033.

Song, B., Scheuner, D., Ron, D., Pennathur, S. & Kaufman, R. J. 2008. Chop deletion reduces oxidative stress, improves beta cell function, and promotes cell survival in multiple mouse models of diabetes. J Clin Invest, 118, 3378–89.

Starr, C. R. & Gorbatyuk, M. S. 2019. Delineating the role of eIF2α in retinal degeneration. Cell Death Dis, 10, 409.

Steinsapir, K. D. & Goldberg, R. A. 2011. Traumatic Optic Neuropathy: An Evolving Understanding. American Journal of Ophthalmology, 151, 928–933.e2.

Sun, D., Chen, X., Gu, G., Wang, J. & Zhang, J. 2017. Potential Roles of Mitochondria-Associated ER Membranes (MAMs) in Traumatic Brain Injury. Cell Mol Neurobiol, 37, 1349–1357.

Tan, H.-P., Guo, Q., Hua, G., Chen, J.-X. & Liang, J.-C. 2018. Inhibition of endoplasmic reticulum stress alleviates secondary injury after traumatic brain injury. Neural regeneration research, 13, 827–836.

Tao, W., Dvoriantchikova, G., Tse, B. C., Pappas, S., Chou, T. H., Tapia, M., Porciatti, V., Ivanov, D., Tse, D. T. & Pelaez, D. 2017. A Novel Mouse Model of Traumatic Optic Neuropathy Using External Ultrasound Energy to Achieve Focal, Indirect Optic Nerve Injury. Sci Rep, 7, 11779.

Taylor Ca, B. J., Breiding Mj, Xu L. 2017. Traumatic Brain Injury–Related Emergency Department Visits, Hospitalizations, and Deaths — United States, 2007 and 2013. MMWR Surveill Summ 2017.

Thaung, C., Arnold, K., Jackson, I. J. & Coffey, P. J. 2002. Presence of visual head tracking differentiates normal sighted from retinal degenerate mice. Neuroscience letters, 325, 21–24.

Torrens, J. N., Hetzer, S. M. & Evanson, N. K. 2023. Brief Oxygen Exposure after Traumatic Brain Injury Hastens Recovery and Promotes Adaptive Chronic Endoplasmic Reticulum Stress Responses. International Journal of Molecular Sciences, 24, 9831.

Ugbode, C., Fort-Aznar, L., Evans, G., Chawla, S. & Sweeney, S. 2019. JNK Signalling Mediates Context-Dependent Responses to Reactive Oxygen Species in Neurons.

Ventura, R. E., Balcer, L. J. & Galetta, S. L. 2014. The neuro-ophthalmology of head trauma. The Lancet Neurology, 13, 1006–1016.

Villegas, R., Martinez, N. W., Lillo, J., Pihan, P., Hernandez, D., Twiss, J. L. & Court, F. A. 2014. Calcium release from intra-axonal endoplasmic reticulum leads to axon degeneration through mitochondrial dysfunction. The Journal of neuroscience: the official journal of the Society for Neuroscience, 34, 7179–7189.

Voeltz, G. K., Rolls, M. M. & Rapoport, T. A. 2002. Structural organization of the endoplasmic reticulum. EMBO reports, 3, 944–950.

Walter, P. & Ron, D. 2011. The unfolded protein response: from stress pathway to homeostatic regulation. Science, 334, 1081–6.

Wang, Z. F., Gao, C., Chen, W., Gao, Y., Wang, H. C., Meng, Y., Luo, C. L., Zhang, M. Y., Chen, G., Chen, X. P., Wang, T. & Tao, L. Y. 2019. Salubrinal offers neuroprotection through suppressing endoplasmic reticulum stress, autophagy and apoptosis in a mouse traumatic brain injury model. Neurobiol Learn Mem, 161, 12–25.

Wek, R. C. 2018. Role of eIF2α Kinases in Translational Control and Adaptation to Cellular Stress. Cold Spring Harb Perspect Biol, 10.

Wenzhu Zhou, Yidan Liang, Weihong Du, Xinyu Liao, Wenqiao Fu, Shanshan Tian, Yongbing Deng & Jiang, X. 2023. ISRIB improves white matter injury following TBI by inhibiting NCOA4-mediated ferritinophagy. Research Square PrePrints.

Wladis, E. J., Aakalu, V. K., Sobel, R. K., Mcculley, T. J., Foster, J. A., Tao, J. P., Freitag, S. K. & Yen, M. T. 2021. Interventions for Indirect Traumatic Optic Neuropathy: A Report by the American Academy of Ophthalmology. Ophthalmology, 128, 928–937.

Yao, Z., Bai, Q. & Wang, G. 2021. Mechanisms of Oxidative Stress and Therapeutic Targets following Intracerebral Hemorrhage. Oxidative Medicine and Cellular Longevity, 2021, 8815441.

Ying, Z., Misra, V. & Verge, V. M. K. 2014. Sensing nerve injury at the axonal ER: Activated Luman/CREB3 serves as a novel axonally synthesized retrograde regeneration signal. Proceedings of the National Academy of Sciences, 111, 16142–16147.

Zhang, P., Hamamura, K., Jiang, C., Zhao, L. & Yokota, H. 2012. Salubrinal promotes healing of surgical wounds in rat femurs. Journal of Bone and Mineral Metabolism, 30, 568–579.

Zito, E. 2015. ERO1: A protein disulfide oxidase and H2O2 producer. Free Radical Biology and Medicine, 83, 299–304.

